# Evolution in response to an abiotic stress shapes species coexistence

**DOI:** 10.1101/2023.05.06.539716

**Authors:** Inês Fragata, Diogo P. Godinho, Leonor R. Rodrigues, Miguel A. Cruz, Flore Zélé, Oscar Godoy, Sara Magalhães

**Affiliations:** cE3c - Center for Ecology, Evolution and Environmental Changes & CHANGE - Global Change and Sustainability Institute, Departamento de Biologia Animal, Faculdade de Ciências, Universidade de Lisboa, Portugal; Institut des Sciences de l’Evolution de Montpellier (ISEM), Université de Montpellier, CNRS, IRD, EPHE, Montpellier, France; Estación Biológica de Doñana (EBD-CSIC), Americo Vespucio 26, E-41092 Sevilla, Spain; Wissenschaftskolleg zu Berlin, Institute for Advanced Study, Berlin, Germany

**Author notes:** Corresponding author: Inês Fragata. co-first (Inês Fragata, Diogo P. Godinho). co-last (Oscar Godoy, Sara Magalhães). Current address: Gulbenkian Institute for Molecular Medicine (GIMM), Oeiras, Portugal.

**Keywords:** Experimental Evolution, Coexistence theory, Competition, Spider mites

## Abstract

Adaptation to abiotic stresses generally relies on traits that are not independent from those affecting species interactions. Still, the impact of such evolutionary processes on coexistence remains elusive. Here, we studied two spider mite species evolving separately on tomato plants that hyper-accumulated cadmium, a stressful environment for herbivores, or on plants without cadmium. Through experimental evolution and structural stability theory, we found that both species coexist in the cadmium environment, but evolution of a single species in cadmium leads to exclusion. However, when both species evolve in cadmium they can coexist. This shift occured due to a simultaneous increase in intra and a decrease in interspecific competition in that environment. These predictions were further confirmed with complementary experiments of population dynamics. Therefore, population shifts to novel environments, even in absence of interspecific competitors, may have unforeseen evolutionary consequences for community composition and the maintenance of species diversity.

## Introduction

Understanding how evolutionary dynamics shape and are shaped by long-term species persistence is key to predicting community composition and biodiversity maintenance. Character displacement in response to the presence of competitors is one of the classical examples of the role of natural selection in shaping trait evolution and species distribution (Brown & Wilson 1956; Lack 1947; Slatkin 1980). Theory predicts that evolving with competitors may lead to changes in competitive interactions (i.e., the negative per-capita effect of one species on another; Abrams & Matsuda 1994; Bernhardt *et al*. 2020; Sakarchi & Germain 2023), which in turn can affect species coexistence (Edwards *et al*. 2018; Germain *et al*. 2018; Vasseur *et al*. 2011; Yamamichi & Letten 2021). In line with these predictions, empirical studies have shown that short term evolution in the presence of competitors can modify competitive traits (Hart *et al*. 2019) and change coexistence patterns (Germain *et al*. 2020; Hiltunen *et al*. 2017; Lankau 2011; Sakarchi & Germain 2023; Zhao *et al*. 2016).

Although the abovementioned studies have demonstrated that short-term evolution is an important force shaping ecological patterns, we may still be under-estimating the potential role of evolution in species coexistence. An important yet overlooked possibility is that traits that affect interactions between species may be selected even in the absence of competitors. For example, species arriving in a vacant environment may rapidly adapt and, potentially monopolize that environment (Nadeau *et al*. 2021). Likewise, prior adaptation to a given environment can lead to a lower relative growth rate of late arriving species (Low-Décarie *et al*. 2011). Moreover, trait evolution in a given environment may affect species community composition in other environments (Fukano et al. 2022; Gallego & Narwani 2022; Limberger & Fussmann 2021). However, to date no study has tested if and how trait evolution in response to an abiotic selection pressure can change intra- and interspecific competition and its impact on species coexistence.

Recent advances of the complementary theoretical frameworks of modern coexistence theory and structural stability (Chesson 2000; Saavedra *et al*. 2017; Yamamichi *et al*. 2022) provide clear mechanistic pathways into how competing species coexist in ecological timescales. In particular, structural stability defines that coexistence is possible when the feasibility domain (defined through species interactions) can accommodate the differences in intrinsic growth rates between species (Godoy *et al*. 2018; Saavedra *et al*. 2017). Combining this framework with experimental evolution allows unraveling whether trait evolution alters the coexistence opportunities or differences in specieś performance, and enhables identifying specific mechanisms by which evolutionary changes affect ecological dynamics.

Here, we assessed how evolution in response to an abiotic stress affects species coexistence by combining two powerful tools: experimental evolution and structural stability, applied to two closely-related spider mite species, *Tetranychus urticae* and *T. evansi*. Spider mites rapidly adapt to novel host plants (Magalhães *et al*. 2007a; Sousa *et al*. 2019; Wybouw *et al*. 2015) and often form host races (Forbes *et al*. 2017; Magalhães *et al*. 2007b). However, they are also found on plants to which they are not adapted, given their passive dispersal (Bitume *et al*. 2011; Fronhofer *et al*. 2014), thus engaging in competitive interactions in less favorable environments. Competitive interactions between *T. urticae* and *T. evansi* have been documented on tomato plants (*Solanum lycopersicum*) (Fragata *et al*. 2022; Sarmento *et al*. 2011). These plants can hyper-accumulate cadmium in their shoots, which strongly reduces spider mite fecundity and survival, producing a strong selective pressure (Godinho *et al*. 2018, 2023, 2024). Here, we studied how adaptation of these two species to an abiotic selection pressure affected the probability of coexistence.

## Material and methods

Details of the maintenance of outbred and experimental evolution populations, common garden procedures and all experiments performed are available in Supplementary Material and Methods.

### Experimental evolution

Experimental evolution populations were initiated by transferring 220 adult mated females of *Tetranychus urticae* or *T. evansi* from outbred populations (Godinho et al. 2020, 2024) to an experimental box containing four tomato leaves and water. We established five replicate populations for each selection regime (Fig. S1A): *T. urticae* or *T. evansi*, exposed to leaves from plants watered either with a 2mM cadmium solution (the cadmium selection regime) or with water (the no-cadmium selection regime). This cadmium concentration is highly detrimental to both spider mite species, (Godinho *et al*. 2018, 2023), a result we recapitulate here (cf.Supplementary Material). Every two weeks (circa one mite generation), 220 adult mated females were transferred to a new box containing four new tomato leaves, ensuring discrete generations. This setup mimics colonization of a new plant, as dispersal is mainly performed by young mated females (Li & Margolies 1993). If the total number of mated females did not reach 220, the transfer was complemented with females taken from the respective T-1 box or from the base population (cf. Godinho et al. 2020, 2024). Replicate 2 of the *T. urticae* cadmium selection regime was not tested because not enough females were available. Prior to the experiments, individuals from all regimes were placed in a common garden of cadmium-free tomato leaves for two generations, to equalize potential maternal effects and synchronize replicates (generation 40 and 42 for the competitive ability and population growth experiments, respectively).

### Empirical estimation of competitive abilities and intrinsic growth rate

To test if evolution on plants with or without cadmium affected the probability of coexistence between the two species in these environments, we estimated the intrinsic growth rates and the strength of intra- and interspecific interactions of mites from each selection regime in each environment (Fig. S1B), following the methodology described in Hart *et al*. (2018). Briefly, after two generations of common garden, we placed one focal female alone or with 1, 3 or 9 females (i.e., competitors) on leaf disks from plants with or without cadmium. The focal females were exposed to competitor females either from the same (intraspecific) or a different (interspecific) selection regime (full factorial design) with matching population replicates. Females were left to oviposit for three days, then killed. Two weeks later, the number of adult females per patch was counted, representing the combined effect of competition on the females and their offspring. Each experimental treatment (i.e. combination of density*selection history of focal and competitor*environment) was replicated 10 times for each replicate population of the four selection regimes, making a total of 1260 replicates divided by six blocks.

### Experimental validation of population dynamics in the presence of interspecific competitors

To test if evolution on plants with or without cadmium affected the population growth rate under competition, we placed six females of the two species from different selection regimes, after two generations of common garden, on plants with or without cadmium (full factorial design, Fig. S1C) and measured the number of adult females produced by each species after two generations. Ten experimental replicates were tested per experimental condition, per replicate population. This data was then compared with model predictions based on parameters estimated from the intrinsic growth rate and competitive abilities (cf. Model validation section below).

### Theoretical estimation of competition and growth parameters

Data collected in the intrinsic growth rate and competitive ability experiment was used to parameterize the Ricker competition model (Fowler 1981; Ricker 1954), used in a previous study with spider mites (Bisschop *et al*. 2022). This model allows incorporating positive interactions (i.e., facilitation, Bowler et al. 2022; Buche et al. 2025; Martyn et al. 2021; Ricker 1954; Stouffer 2022), which, based on initial data scrutiny, were likely to occur in our system. The Ricker model is described by the following equation:

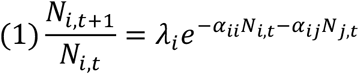

where *N_i,t+1_* is the number of individuals of species *i* in the next generation, 𝜆_!_the intrinsic growth rate of species *i* in absence of competitors, 𝛼*_ii_* and 𝛼*_i_*_)_ the strength of intra- and interspecific competition, respectively, and *N_i,t_*, *N_j,t_* the number of competitors of species *i* and *j*, respectively, in the current generation. We used the “cxr” package (García-Callejas *et al*. 2020) in R to estimate 𝜆*_i_, α*_ii_ and *α_ij_* for each replicate population separately and with all replicates of the same selection regime pooled (to increase statistical power). To include the filtering effect of the cadmium environment and because the “cxr” package does not allow the inclusion of zeros, we transformed our data by summing one to each of our datapoints (as the logarithm of one is zero). The initial parameters to be inputed in the “cxr” package were selected based on the mean likelihood within each environment after performing an exploration of the likelihood surface of the models (details in Supplementary Methods). Model fitting assessment was done by visual inspection (availabe in the git repository) and by comparing the Euclidian distance between observed and predicted values (Fig. S2). To assess the relative impact of the different parameters on population growth, we simulated offspring production of ten females using only the intrinsic growth rate, the joint effects of intrinsic growth rate and of intraspecific competition or the joint effects of those traits plus interspecific competition. In general, confidence intervals were calculated using the lower and upper estimates obtained from the 2000 bootstrap iterations from the cxr package (García-Callejas *et al*. 2020).

### Structural stability approach to predict coexistence

Estimates obtained from the Ricker model (Fig. S3, S4, S5, S6) showed widespread weak positive interspecific interactions (i.e., facilitation), which are not accounted for in common approaches to study species coexistence (e.g. Chesson 2000). Therefore, we used the structural stability framework (Allen-Perkins *et al*. 2023; Saavedra *et al*. 2017, 2020) to predict the outcomes of species interactions. This approach utilizes the strength and sign of species interactions to estimate the size of the feasibility domain, and allows incorporating stochasticity and interactions other than direct competition (Rohr *et al*. 2014). Coexistence is possible if the vector containing the intrinsic growth rate of both species falls within the feasibility domain.

In principle, the latter can include both negative and positive intrinsic growth rates. However, we restricted coexistence predictions to positive values (i.e. bounded by the vectors 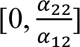 and 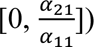 to match information from our experimental system, as species can either have zero or positive growth rates (Song *et al*. 2018). Finally, to account for uncertainty in the estimation of model parameters, we estimated the feasibility domain bound by the upper (lower) 95% of the vector 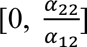 and the lower (upper) boundaries of the 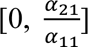 to obtain the widest (narrowest) conditions under which species are predicted to coexist.

To measure each species’ vulnerability to exclusion, we tested how resistant coexistence is to perturbations. For that we estimated the species exclusion distance (following Allen-Perkins *et al*. 2023; Medeiros *et al*. 2021), which corresponds to the minimal distance between the vector of intrinsic growth rates 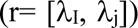 and the edges of the feasibility domain (vectors corresponding to either 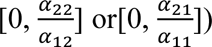 as a proxy for how strong a perturbation must be to change the coexistence outcome. To calculate the distance between the normalized vectors of intrinsic growth rates and the edges of the feasibility domain, we applied the following formula, (Allen-Perkins *et al*. 2023):

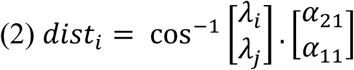

Uncertainty in the estimated distance was accounted for by estimating the distance between the normalized vectors of intrinsic growth and the edges of the smallest or largest feasibility cones. The largest and smallest feasibility cones were obtained by using the lower and upper parameter estimates, respectively, to encompass potential trait variation underlying observed parameter estimates.

### Model validation

We validated our approach by comparing the projections of abundances in the short term, obtained from our model, with the empirical results of a population growth experiment after two generations. For that, we used the Ricker model to predict the relative abundance of each species after two generations, starting with the initial conditions of the experiment. Then, we performed a general linear regression with a gamma error distribution to estimate the correlation between the mean observed proportion of T. evansi females (Number of *T. evansi* females/Number of females) in all replicates of all treatments with the model proportion estimates obtained for each replicate.

### Statistical analyses

To test how evolution on plants with cadmium affected population growth and competition in the cadmium environment, we used general linear models with a gamma distribution for the intrinsic growth rate, and with a normal distribution for intra- and interspecific competition. All models were applied separately for both species, and normality and dispersion of the residuals was inspected for all models. We included the parameters estimated with the cxr package for each replicate as dependent variables and selection regime as a fixed factor (with two levels, cadmium and no-cadmium). For the strength of interspecific competition, the model included the selection regime of the focal and competitor species, and their interaction. Whenever the interaction term was significant, we applied contrasts for all combinations of selection regimes. Similar models were applied to test the impact of evolution in cadmium on the performance in the no-cadmium environment. A similar approach was used to test the impact of cadmium on the performance of the no-cadmium selection regimes (but with factor Environment as fixed factor). Finally, we estimated the likelihood of finding differences between environments/selection regimes using 10000 bootstrap iterations. For that, we randomized (with replacement) the parameter estimates within environments/selection regimes (depending on the model) and applied the same models as above. Then, we estimated the probability of randomly obtaining a significant p-value (below or equal to 0.05) with our data set by computing the number of times a significant result was obtained.

All analyses were done using the package “glmmTMB” (Brooks *et al*. 2017) in R 4.2.1 version (R Core Team 2022). Contrasts were performed using the “emmeans” package (Lenth 2024), analyses of residuals were done for each model using the DHARMa package (Hartig 2022), and graphical representation was done using “ggplot2” (Wickham 2016). All data and scripts are available in the repository https://figshare.com/s/f001d9f699a4027d7b62.

## Results

### Evolution in cadmium changes population growth and the interactions within and between species

In the cadmium environment, the intrinsic growth rate of both species evolving in that environment for 40 generations was higher than that of populations evolving on plants without cadmium, although this value was only significant for *Tetranychus urticae* (*T. urticae*: χ^2^_1,6_=6.012, P-value=0.014, *T. evansi*: χ^2^_1,7_= 2.796, P-value=0.095, Fig 1, Fig. S7, Table S1, S2). In addition, the strength of intraspecific competition increased for *T. evansi* (χ^2^_1,7_ = 4.957, P-value=0.0259, Fig S3A, Table S1, S2), but not for *T. urticae* (χ^2^_1,6_ =0.495, P-value=0.4818, Fig S3B, Table S1, S2). This resulted in a decreased predicted number of offspring produced by *T. evansi* in competition (relative to when growing alone) but only when *T. evansi* evolved in the cadmium environment (Fig. 1A, λ vs λ+α_ii_ for the yellow and blue vs the green and red dot). Moreover, evolving on plants with cadmium made *T. evansi* more sensitive to competition with *T. urticae* from the no-cadmium selection regime (Fig. S4A, Table S3A, S3B, contrasts: T ratio_1,13_= −3.150, P-value=0.0340), reducing the predicted number of offspring produced after one generation of competition (Fig. 1A, λ+α_ii_+α_ij_, comparing yellow and red dots). Conversely, evolution in cadmium did not affect the sensitivity to interspecific competition in *T. urticae* (Fig. S4B, Table S3A). We observed weak positive interactions within and between species in several replicates (i.e., negative alpha values) across selection regimes (Fig S3, S4), which reinforces the added value of using the structural stability framework.

**Figure 1-.**
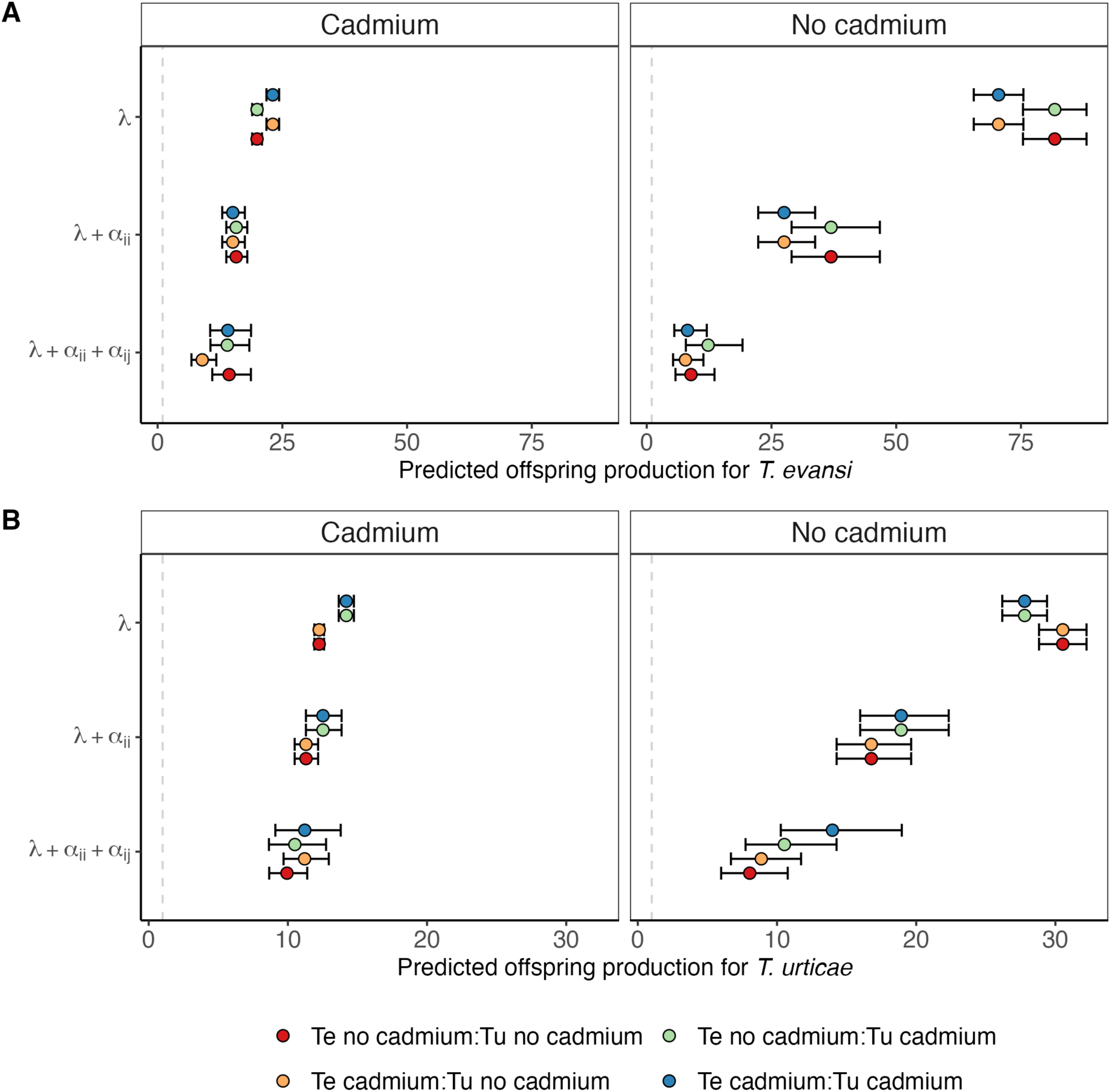
Predicted number of female offspring after one generation for A) *T. evansi* (Te) and B) *T. urticae* (Tu), to estimate the relative effect of competition on the growth rate of the different populations. To represent the cumulative impact of each parameter on offspring production, predictions were done using only the intrinsic growth rate (𝜆), the intrinsic growth rate plus intraspecific competition (𝜆 + 𝛼*_ii_*) and the latter plus interspecific competition (𝜆 + 𝛼*_ii_* + 𝛼*_i_*_)_) in an environment with or without cadmium (left and right panels, respectively). Note that, since two of the four treatments share the same focal selection regime (e.g. for *T. urticae* the orange and red dots correspond to the focal control regime “Tu no-cadmium”), two colours are duplicates for the metrics without interspecific competition (𝑖. 𝑒., 𝜆 and 𝜆 + 𝛼*_ii_*). Starting conditions for simulations: 10 individuals of each species. Treatments correspond to combinations of *T. evansi* (Te) and *T. urticae* (Tu) selection regimes (cf. colour code). Note the different scales on the X-axis between the two species (A vs B). Confidence intervals for each prediction were estimated using the lower and upper estimates of the intrinsic growth rate and intra and interspecific competition indexes obtained from the model.

In the no-cadmium environment, no significant differences in the intrinsic growth rate (Fig. S8, T. *evansi*: χ^2^_1,7_=0.015, P-value=0.969; T. *urticae*: χ^2^_1,6_= 0.4988, P-value=0.48, Table S1, S2), intraspecific competition (Fig. S5, *T. evansi:* χ^2^_1,7_= 2.5448, P-value=0.1107; *T. urticae*: χ^2^_1,6_= 1.3944, P-value=0.2377, Table S1, S2) or interspecific competition (Fig. S6, Table S1, S2, S4) were observed between populations evolving in that environment and those evolving in the environment with cadmium. Weak intra- and interspecific facilitation was found for several replicates across selection regimes (Figs. S5, S6), as in the cadmium environment.

### Evolution of both competitors increases the probability of long-term coexistence in the cadmium environment

In the cadmium environment, coexistence was more likely when either none of the species evolved in cadmium or when both species had evolved in that environment (Fig. 2). Coexistence when both species evolved in the no cadmium environment was promoted by a decrease in the growth rate and in the strength of intra and interspecific competition for the two species (Table S5). These changes led to an increase in the size of the feasibility domain (Fig. 2), except when considering the narrower parameter estimates (Fig 2, light blue region). Coexistence when both species evolved in cadmium was due to a larger increase in the size of the feasibility domain via increased intraspecific competition for the better competitor (*T. evansi*, Fig S3). The prediction of coexistence of both species after evolution in cadmium was robust to parameter variation because it holds even after taking uncertainty (95% CI) into account when estimating the feasibility domain (Fig 2, light blue region), and the differences of species intrinsic growth rates (Fig. 3, S9).

**Figure 2-.**
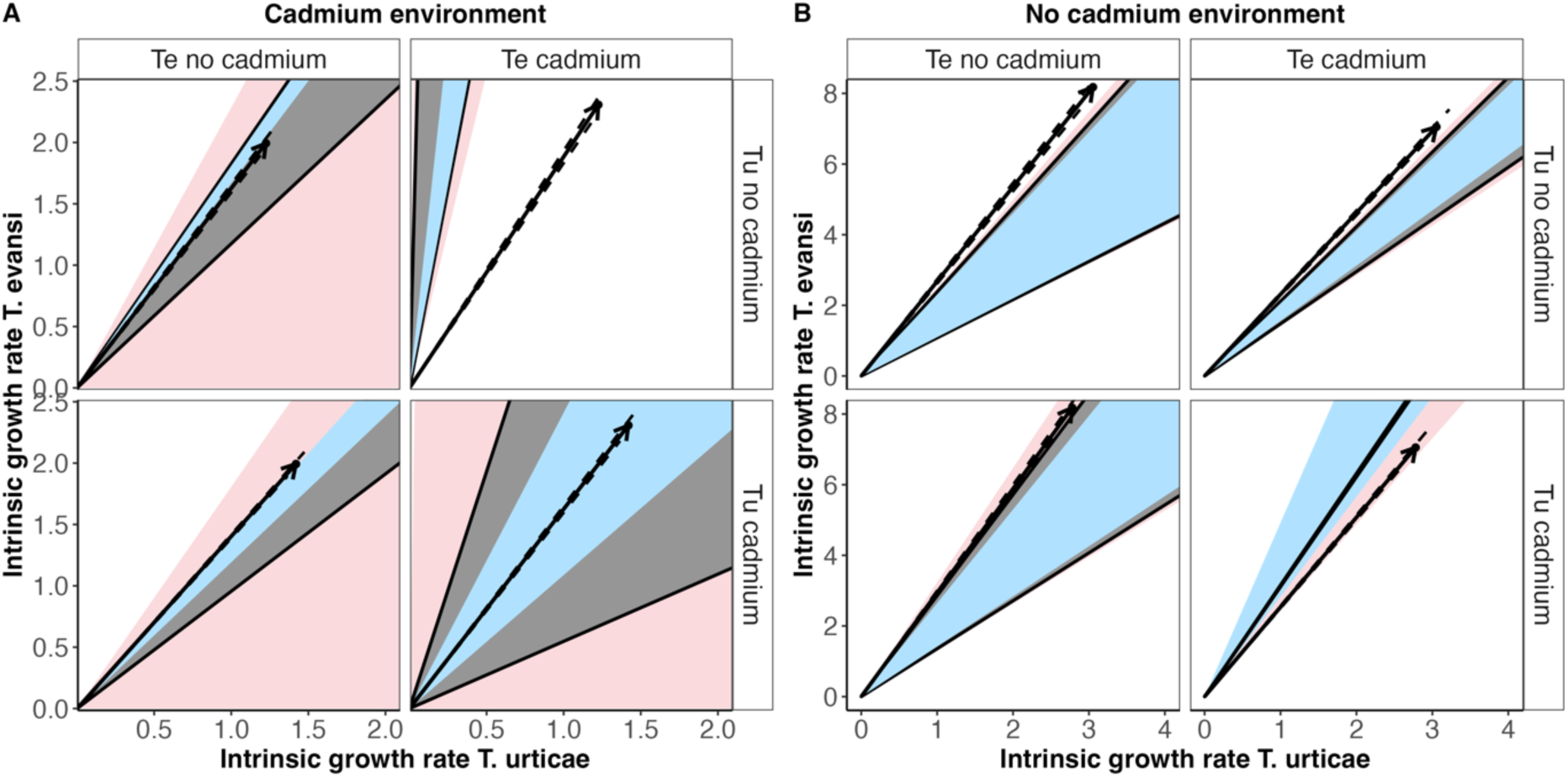
Feasibility domain predicted for enviromments with (A) or without (B) cadmium for the different combinations of cadmium and no-cadmium selection regimes in which *T. urticae* (Tu) or *T. evansi* (Te) have evolved. The x and y axis correspond to the positive intrinsic growth rate for *T. urticae* and *T. evansi*, respectively. The arrow represents the vector of the intrinsic growth for each of the two species. The black lines delimiting the grey cone indicate the area under which the isoclines cross at positive abundances (i.e., the feasibility domain, in which coexistence is possible). Dashed lines surrounding the arrows represent the 95% confidence interval of the intrinsic growth rate. Arrows that fall outside of the cone indicate that Te excludes Tu, except in the upper-right panel of figure A (Te cadmium-Tu no-cadmium) and lower-right panel of figure B (Te cadmium-Tu cadmium), in which Tu excludes Te. The dark grey region indicates the feasibility domain obtained from the ratio of intra and interspecific competitive abilities estimated with data from all replicates pooled (see methods). This region has been evaluated only under the conditions in which both species show positive intrinsic growth rates. The light blue and light red regions delimitate the smallest and largest feasibility domains obtained from the lower and upper 95% confidence interval of the competitive ability parameters (the light blue region is always contained within the feasibility cone in grey). White regions denote the unfeasibility domain (region for exclusion). Confidence intervals were obtained from the upper and lower parameters estimates from the data of all experimental replicates pooled.

**Figure 3-.**
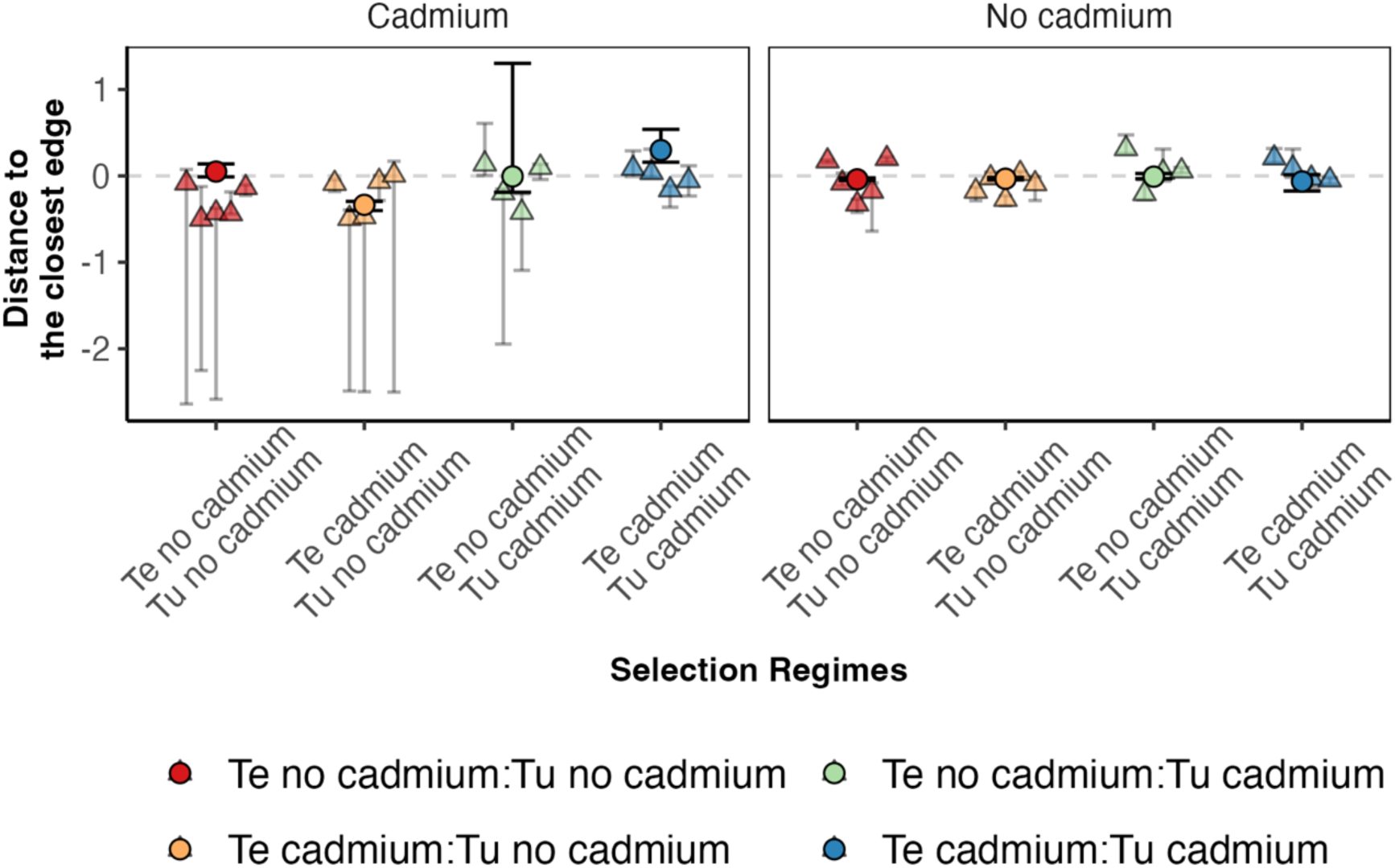
Distance between the vector of intrinsic growth rates and the closest edge of the feasibility domain in environments with (left panel) or without (right panel) cadmium for the different combinations of no-cadmium or cadmium selection regimes for *T. evansi* (Te) and for *T. urticae* (Tu) (cf. colour code). Positive distances indicate that the vector of growth rates is inside the feasibility domain (i.e., coexistence is possible), and negative distances indicate that the vector is outside the feasibility domain (i.e., exclusion is predicted). Circles correspond to the distance calculated with all data pooled together, and triangles to distances calculated per replicate. Confidence intervals describe the difference between the vector of intrinsic growth rates and the edges of the largest or smallest feasibility domain, which were obtained from the lower and upper estimates of the parameters as described in Figure 2. The pooled data estimates were obtainedfrom data from all experimental evolution replicates to increase statistical power and to account for the variation between replicates.

In the cadmium environment, coexistence was not possible when only one species evolved in that environment (Figs 2, 3). In fact, we predicted competitive exclusion for whichever species evolved in cadmium (Figs. 2, 3, S9). The exclusion of *T. evansi* after evolving in cadmium can be explained by two factors: increased intraspecific competition (Fig S3) and increased sensitivity to competition (i.e., stronger response to competition) from *T. urticae* of the no-cadmium selection regime (Fig S4). In the reverse case, the exclusion of *T. urticae* that evolved in cadmium (Fig. 4), can be explained by an assymetric effect of interspecific competition. *T. urticae* barely affecting *T. evansi* from the no-cadmium selection regime, but the latter strongly affecting *T. urticae* (Fig. S4). Again, these predictions, were robust to variation in parameter estimation when accounting for lower and upper parameter estimates.

**Figure 4-.**
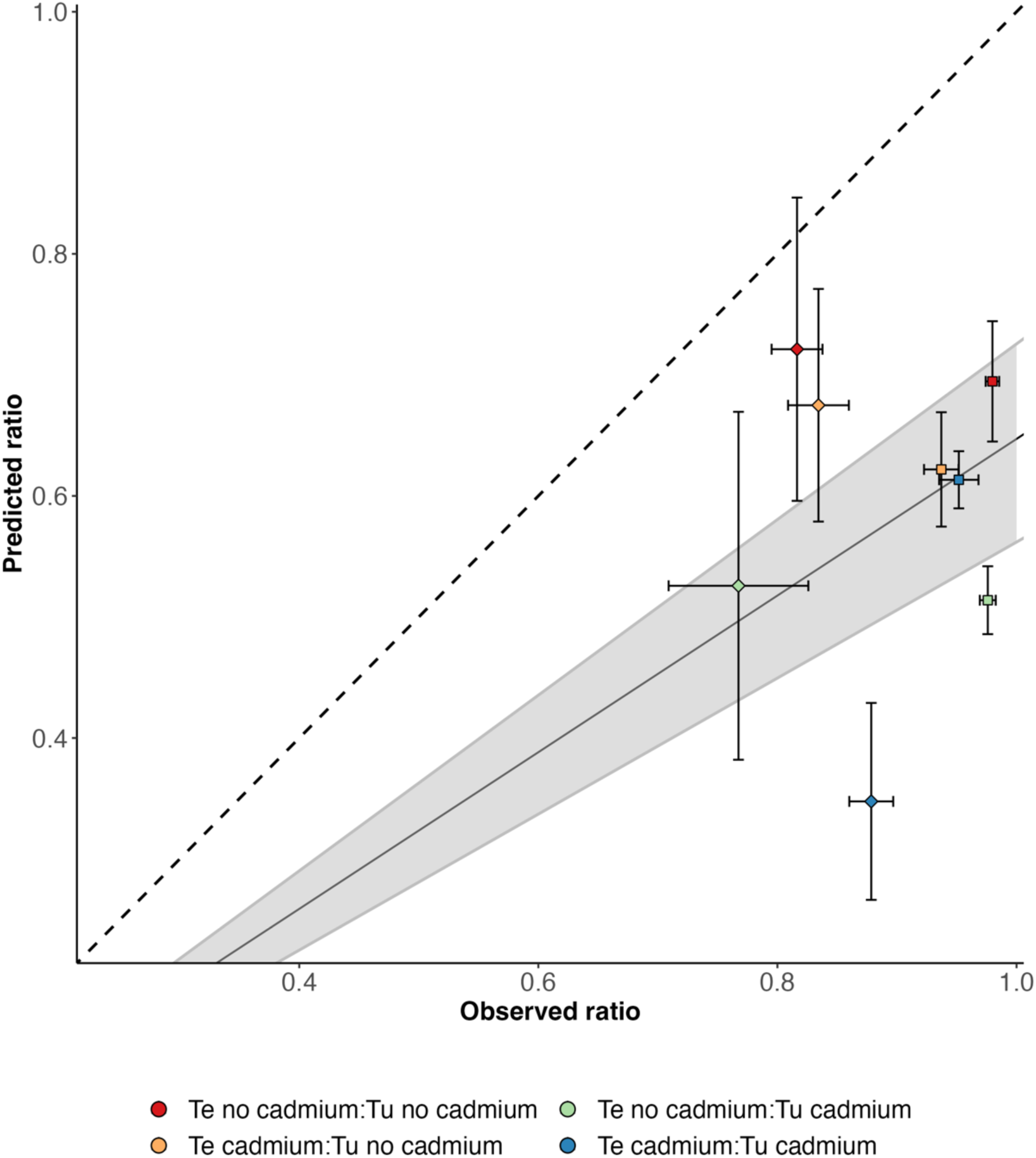
Correlation between predicted and observed relative abundance of *T. evansi* (Number of *T. evansi* females/Number of females of both species) after 2 generations in an independent population growth experiment in which the two species are released at equal densities. Squares indicate data from the cadmium environment and diamonds data from the no-cadmium environment. The full line represents the slope obtained from a general linear model (see methods) and the shaded area corresponds to the confidence interval for the slope obtained from the model. Treatments correspond to combinations of selection regimes to which *T. evansi* (Te) and *T. urticae* (Tu) were exposed (cf. colour code). Predicted error bars were calculated based on the upper and lower parameter estimates of all data pooled and observed error bars were obtained from the standard error of the replicate populations. Dashed black line indicates 1:1 ratio between predicted and observed values.

In the no-cadmium environment, exclusion was generally predicted, with *T. urticae* being excluded most for most scenarios, except when both species evolved in cadmium (Fig 2). However, the minimum distance to the edges of the feasibility domain was short in all cases (Fig. 3, S9), especially when considering those involving lower confidence intervals, indicating that small perturbations may shift the prediction from coexistence to exclusion and vice versa. Importantly, there was high heterogeneity among replicates but little effect of the evolutionary history of both species (Fig. 3). Thus, evolving in the cadmium environment did not affect the range of opportunities for species to coexist in the no-cadmium environment (Fig. 2, S9).

### Simulated competitive dynamics partially recover empirical patterns

To validate our approach, we compared theoretical predictions with a two-generation population dynamic experiment with the two species from all combinations of selection regimes. (Fig. S1). Results show that we consistently underestimated the proportion of *T. evansi* in the populations. Still, the model was able to recapture the tendency observed in the data (Fig. 4, slope pooled replicates: 0.647, P-value<0.001), despite some heterogeneity between replicate populations, especially in the no-cadmium environment (slope with replicates: 0.642, P-value<0.001, Table S6).

## Discussion

We used a combination of experimental evolution and structural stability theory to predict the impact of evolution of two spider mite species in response to an abiotic selection pressure (cadmium) on species persistence, in an environment with or without cadmium. To obtain theoretical predictions, we quantified the intrinsic growth rate of each species and the strength of their intra- and interspecific competitive interactions across different environmental and selection scenarios. We find that cadmium equalized fitness differences between species, decreasing growth rate and competitive interaction (within and between species). Interestingly, evolutionary history had little effect on coexistence patterns in the environment without cadmium. However, the independent evolution of both species in presence of cadmium led to differential changes in intrinsic growth rates (which equalized fitness between species), coupled with an increase in intraspecific competition in the superior competitor (which led to increased structural niche differences), and therefore to an increased range of fitness differences compatible with persistence of both species in the cadmium environment. However, this only occurred when both species evolved in cadmium. In sum, we show that evolution in response to an abiotic selection pressure can change interactions between and within populations and affect the probability of coexistence, even without a direct selection pressure posed by the presence of competitors as showed previously (Germain *et al*. 2020; Sakarchi & Germain 2023; Vasseur *et al*. 2011; Zhao *et al*. 2016).

Adaptation to cadmium occurred only for the worst competitor (*T. urticae*), as evidenced by the increase in the intrinsic growth rate in the cadmium environment of populations from the cadmium selection regime, when compared to the no-cadmium regime. Previously we reported that *T. evansi* populations did not shown signs of adaptation at the 33rd generation of evolution (Godinho et al. 2024), but showed a reduced performance in the no cadmium environment. Here we observe that populations evolving in cadmium show a marginally significant increase in the intrinsic growth rate in the cadmium environment, suggesting that adaptation may still be in process. However, we do not recover the loss of performance in the no cadmium environment, suggesting that extending time for evolution to operate led to reduced costs of evolving in the cadmium environment. This pattern of slow increase in growth rate, together with the lack of initial genetic variation for traits associated to performance in cadmium, is compatible with the existence of cryptic genetic variation that is released upon evolution in the cadmium environment, for example via newly formed epistatic interactions (Paaby & Rockman 2014). As the traits we are measuring are likely polygenic, this leads to more allele combinations, thereby increasing the likelihood of such cryptic genetic variation to arise (Chandler 2010; Lauter & Doebley 2002). Still, adaptation may be too slow to enable the establishment of mites in cadmium-contaminated sites. Indeed, assuming similar generation time in the lab and in the field, a tomato growing season roughly corresponds to 15 generations for spider mites, which is not sufficient to lead to genetic changes allowing adaptation to cadmium.

Evolution of *T. evansi* in cadmium also led to stronger sensitivity to intraspecific competition in that environment, as compared to mites evolving on plants without cadmium. Evidence that intraspecific competitive ability may evolve during adaptation to a novel abiotic environment has been accumulating (Bernhardt *et al*. 2020; Bono *et al*. 2015; Limberger & Fussmann 2021). There are many reasons to expect such evolution (Siepielski *et al*. 2016). For example, individuals may become more efficient at extracting resources when these are limiting (Bernhardt *et al*. 2020), or higher population growth may lead to higher densities being reached earlier, thus increasing intraspecific competition. This effect is expected to be stronger for *T. evansi*, given that its intrinsic growth rate is higher than that of its competitor. Measuring intraspecific competition should thus be mainstreamed in experimental evolution studies, which typically measure only individual life-history traits such as fecundity and survival (Kawecki *et al*. 2012).

An overlooked logical follow-up of our observations is that adaptation to abiotic selection pressures may affect interspecific interactions as well. Here, we shed light on this understudied hypothesis by documenting that, when spider mites evolved in cadmium, they affected their competitors less or equally than mites that evolved on plants without cadmium. This counter-intuitive result may be due to a trade-off between inter and intraspecific competition, given that *T. evansi* shows increased sensitivity to intraspecific competition and reduced sensitivity to interspecific competition on plants with cadmium upon evolution in that environment. Such trade-off has been assumed in theoretical models (Vasseur *et al*. 2011), and experimentally demonstrated in *Brassica nigra* (Lankau 2008; Lankau & Strauss 2007). Evolution of interspecific competitive ability has been also observed when competing species coevolve (Fukano *et al*. 2022; Germain *et al*. 2020; Hart *et al*. 2019; Sakarchi & Germain 2023). The fact that we recover such patterns even in the absence of a competitor in the environment implies both that causality needs to be scrutinized in eco-evolutionary studies and that evolution has consequences for community structure that are much more far reaching than initially thought.

Empirical studies have shown that adaptation to an abiotic selection pressure can change species interactions (Bach *et al*. 2018; Fukano *et al*. 2022; Limberger & Fussmann 2021) but none have addressed how it affects species coexistence. In fact, the only studies applying coexistence theory to evolutionary data concern adaptation to the presence of a competitor (Germain *et al*. 2020; Hart *et al*. 2019; Zhao *et al*. 2016). Here, we found that, in the cadmium environment, species are predicted to coexist either when neither evolved in the cadmium environment or when both competitors evolved separately in cadmium. Given the mosaic structure of spider mite populations in the field (Magalhães *et al*. 2007b), their rapid adaptation to novel host plants (Sousa *et al*. 2019) and the passive nature of their dispersal (Smitley’ & Kennedy 1988) this scenario is quite plausible.

The increase we observed in the range of the feasibility domain after evolution in cadmium was not due to a large change of a specific parameter but rather a small but full reorganization of how species grow and interact with each other. Namely, coexistence was due to a combination of a stronger self-limitation of the superior competitor with a reduced negative competitive effect on the inferior competitor. Although the ultimate mechanisms of these changes are unclear, we speculate that evolution improved the ability of spider mites to feed on tomato plants that accumulated cadmium, likely by increasing resource uptake, which in turn decreased resources available to others (thus increasing intraspecific competition). This positive covariance between an increase in fitness and an in self-limitation is congruent with theoretical predictions (Carroll *et al*. 2011), previously reported in other systems (e.g. Angert et al. 2009). Additionally, the observed reduction in interspecific competition could be due to plant defence suppression, documented in these populations (Fragata *et al*. 2022). Such suppression may make resources more available to both species. Another possible mechanism affecting competition in our system is resource heterogeneity among leaves, which, together with differences in arrival, have been shown to shape coexistence between these two species (Fragata *et al*. 2022). Studies addressing how evolution in the presence of a competitor affected species coexistence have found that coexistence is maintained when species coevolve, albeit by mechanisms different than those operating in the absence of such evolution (Germain *et al*. 2020; Hart *et al*. 2019; Zhao *et al*. 2016). Given that habitats are increasingly fragmented in space and changing in time, organisms are often exposed to different abiotic challenges. We show that they may rapidly adapt to these changes, fostering their ability to persist in those environments, both directly via coping with abiotic challenges, and indirectly, via the effect of such adaptation on species interactions. Therefore, adaptation to an abiotic selection pressure may affect community composition to an unprecedent degree, higher than direct adaptation to the presence of competitors.

We also tested the accuracy of the theoretical predictions, by comparing model predictions based on parameters estimated from competition experiments with a short-term population dynamics experiment. Despite the fact that our model consistently underpredicted the proportion of *T. evansi* in the population (as the slope was lower than 1), we still obtain a good overlap between predicted and observed relative abundances of the two species. Some mismatch was found for the no cadmium environment when both species had similar history and for the cadmium environment when only *T. urticae* evolved in that environment. This is probably due to stochasticity, as the experiment started with a small number of individuals. We also obtain large confidence intervals for the competitive interactions estimates, which may be due to low statistical power (especially when estimating per replicate population), but can also be caused by non constant effect of competitive interactions that is not captured by our current approach. In the future, it will be important to expand the structural stability approach to include different types of responses to competiton. Still, the strong correlation between observed and predicted values suggests that our model parameterization is robust to changes in the number of individuals, resource and space availability, indicating that the combination of theoretical modelling and experimental estimation of competitive interactions has a high predictive power.

Another strength of our study is that, unlike previous studies, we applied a full factorial design, in which we explore how different combinations of evolutionary histories of both competitors affect the probability of coexistence. In fact, when competition occurred between one species that evolved without cadmium and one that evolved in cadmium, the latter was more likely to be excluded in the cadmium environment. This unintuittive outcome may be explained by a combination of increased self-regulation (i.e. increased intraspecific competition) by the cadmium-evolved population and the weaker interspecific competition exerted by the cadmium populations. Hence, community assembly does not follow a linear path, being contingent on the evolutionary history of the two species. This suggests that asynchronies in the arrival to cadmium-contaminated sites may lead to species exclusion, due to evolution of the species first colonizing that environment. Coexistence is only possible when the two species have either both evolved in the no cadmium environment or both evolved in the cadmium environment. The latter can be achieved either if they arrive simultaneously to a site with cadmium, then adapt at a similar pace, or if they are already adapted upon arrival. Hence, as previously shown for short-term differences in arrival time (Fragata *et al*. 2022), we here show that historical contingencies affect species coexistence also via their effect on evolution.

As found in studies with no evolution, we found that interactions are specific to a given environment (Grainger *et al*. 2019; Granjel *et al*. 2023; Matías *et al*. 2018; Wainwright *et al*. 2019). In general, our results align with previous work, not considering evolution, showing that stable coexistence is fostered in stressful environments via a reduction in growth rate that equalizes fitness differences, and by a shift from interspecific to intraspecific competition, which increases the range of coexistence opportunies (i.e. the size of the feasibility domain) (Grainger *et al*. 2019; Granjel *et al*. 2023; Matías *et al*. 2018; Song *et al*. 2020b; Wainwright *et al*. 2019), and stabilize the population dynamics of interacting species. However, our study tempers this statement by the finding that this is only true when there is a match in the evolutionary history of both species in cadmium (i.e. both evolved in cadmium or evolved in the no cadmium environment). Moreover, the use of the structural stability framework allows incorporating sensitivity to changes in environmental conditions in these coexistence predictions (Allen-Perkins *et al*. 2023; Song *et al*. 2020a; Tabi *et al*. 2020). In the environment without cadmium, we found a small distance to the edge (i.e., low robustness) across all selection regimes, suggesting that changes in competitive outcomes are likely to occur due to stochastic events. Instead, in the cadmium environment, the distance to the edge was higher in all cases, suggesting that communities in that environment are more long-lasting. Thus, our results suggest that communities in cadmium-free environments will be modulated by small environmental changes, whereas those in environments with cadmium will be more shaped by evolution.

Our study highlights the added value of combining experimental evolution and controlled experiments with ecological theory, by providing novel insights of how species adaptation to an abiotic stressor affects their ability to coexist. This work provides significant advances in both evolutionary and ecological fields. On the one hand, it shows that traits such as the strength of intra- and interspecific competition should be incorporated in the characterization of species adaptation to novel environments. This is particularly true in experimental evolution studies, which generally quantify evolution by measuring classical life-history traits of single individuals (Garland & Rose 2009; Kawecki *et al*. 2012), thus ignoring traits associated to evolving in the presence of others (Chippindale *et al*. 2003). Conversely, ecological studies benefit from incorporating past evolutionary history, as such approach has the capacity to refine our understanding of how species interact and coexist (Leibold *et al*. 2019; Urban & De Meester 2009; Wittmann & Fukami 2018; Yamamichi *et al*. 2020). Thus, we highlight the need to combine ecological and evolutionary perspectives and methodologies to understand community composition.

## Acknowledgments

This work was financed by an ERC (European Research Council) consolidator grant COMPCON, GA 725419 attributed to SM, an ERC starting grant DYNAMICTRIO, GA 101042392 attributed to IF, and by FCT (Fundação para Ciência e Tecnologia) with the Junior researcher contract (CEECIND/02616/2018) attributed to IF. OG acknowledges financial support provided by the Spanish Ministry of Economy and Competitiveness (MINECO) and by the European Social Fund through the UCCO project (CNS2023-144337). This is contribution ISEM-2023-XXX of the Institute of Evolutionary Science of Montpellier (ISEM). The authors acknowledge stimulating discussion with all members of the Mite Squad and Raul Costa-Pereira, which have significantly improved the experimental design and interpretation of the results, Alexandre Blanckaert for his help with code troubleshooting, and Marta Artal and Liesbeth de Jong for invaluable infrastructure for meetings in Sevilla. For the purpose of Open Access, a CC-BY 4.0 public copyright licence has been applied by the authors to the present document and will be applied to all subsequent versions up to the Author Accepted Manuscript arising from this submission.

## Data accessibility statement

Data and scripts are available in figshare in the link https://figshare.com/s/f001d9f699a4027d7b62, with the doi: 10.6084/m9.figshare.28369472

## Authorship Statement

IF, OG and SM conceived the original idea. IF, DG, OG and SM designed the experiments. IF, DG, LRR and MC performed the experiments and collected the data. IF performed the modelling work and analysed data, with advice from FZ and OG. IF produced the figures, with help from LRR. IF, SM and OG wrote the first draft of the manuscript, and all authors contributed substantially to revisions. SM supervised DG, IF, LRR and MC, FZ supervised MC, OG supervised IF. SM provided the financial support for the experiments, IF and OG for the meetings to analyze the data and write the paper.

