## Supplementary material for "Evolution in response to an abiotic stress shapes species coexistence": SI

#### Supplementary Material and methods

##### General populations maintenance conditions

Outbred populations of each species were maintained using two 5-week-old potted tomato plants placed inside a plastic box, with ventilation. All plants used had at least five fully expanded leaves. Each week a new plant was added and leaves from the discarded plant were added on top of the new plant to allow migration to the new host. Plants used in all experiments and in the experimental evolution were watered twice a week with 100 mL of tap water for a duration of two weeks, then transplanted to new pots and watered each week once with 100mL tap water and another two times with either 100mL of distilled water or 100mL of 2mM cadmium chloride solution during another three weeks. This cadmium concentration has been previously shown to be highly detrimental for both spider-mite species (Godinho *et al.* 2018, 2023). We recover this deleterious effect in our populations (see below).

##### Experimental evolution – Establishment and maintenance

Outbred populations, formed via controlled, one-on-one crosses of 400 individuals (200 females and 200 males) of each of three or four field populations of *T. urticae* and *T. evansi* spider mites, respectively, paired with each other population of the same species. (Godinho *et al.* 2020), were used to initiate experimentally evolving populations. The latter were created by transferring 220 adult females of each species from the outbred populations to an experimental box (size: 26 x 18 x 13.5 cm) covered with a lid with a gauze-covered opening to allow air circulation. The box contained four tomato leaves with their stems in a small pot with water (two leaves per pot, two pots per box), replenished every week. To minimize differences among plants and leaves, each box contained the second, third, fourth and fifth leaves to develop (starting from the cotyledons), taken from a different plant. We established five populations for each of four selection regimes: *T. urticae* or *T. evansi*, exposed to leaves from plants watered either with a 2mM cadmium solution (the cadmium selection regime) or with water only (the no-cadmium selection regime). Every mite generation (i.e. two weeks) 220 adult mated females were transferred by aspiration to a new experimental box containing four new tomato leaves (i.e., populations were maintained in discrete generations). The remaining mites were transferred to a new box with two tomato leaves to create a T-1 box. These mites were used as a backup when less than 220 mated females were found in the experimental box. Females from the base population were only added if when using the T and T-1 boxes, the number of individuals transferred did not reach 220 mated females (cf. (Godinho *et al.* 2020) for details). All experimental evolution populations and experiments were performed in a climatic room under controlled conditions (25:20 ºC, 65% of humidity, light:dark = 16:8).

**Experimental evolution – Common-garden**

Two generations of common garden were performed for all populations prior to each experiment. For that, 200 females from each population were transferred to new boxes (100 per box) with two tomato leaves, except for replicate 2 of *T. urticae* evolving on leaves with cadmium, which did not produce enough individuals and was thus not used in subsequent experiments. After three weeks, six cohorts of 50 adult females were transferred to petri dishes with cotton soaked in water and two tomato leaflets. Females were left to oviposit for four days, then killed. Two weeks later, the adult female offspring was collected and used in the experiment. Experiments were performed at generation 40 (empirical estimation of competitive ability and intrinsic growth rate) and 42 (estimation of the growth rate of populations in cages with interspecific competitors).

**Empirical estimation of competitive abilities and intrinsic growth rate**

To quantify changes in growth rate with different competition scenarios and estimate the parameters to predict coexistence outcomes, we followed the methodology described in Hart et al. (2018). For that, we cut leaf discs (18mm diameter) from tomato plants with or without cadmium and placed one female alone or with 1, 3 or 9 females (i.e., competitors), either from the same or a different species, to estimate intraspecific or interspecific competition, respectively. In intraspecific competition, the competitor females were from the same selection regime (cadmium or no-cadmium) and replicate population as the focal female. In interspecific competition, the focal females were exposed to females from either the same or a different selection regimes (full factorial design). For all tests we used individuals from populations with the same replicate number, as they were transferred in the same day. The intrinsic growth rate was estimated from leaf discs with single females.

Leaf discs were placed on water-saturated cotton inside a square petri dish (size: 12.6 x 12.6 x 1.6 cm, with a gauze covered top to allow for ventilation), with 16 or 12 discs (for intra and interspecific assays, respectively). Each petri dish corresponded to one environment (leaf discs from plants exposed to cadmium or not) and one treatment (a combination of the selection regimes of the focal female and of the competitors – in the case of intraspecific competition these are the same), with each density of competitors being allocated to one row within the petri dish (i.e., each petri dish had four replicates of all densities). After three days, the state of the focal and competitor females was scored (alive, dead, drowned or missing) and females were removed. As the eggs of the two species cannot be distinguished, we counted the number of alive or dead adult daughters per patch instead. For each of the five experimental evolution replicates we performed ten replicates per intra- or intraspecific treatment. The experiment was done in six blocks, the first three with replicate populations one to three and the last with replicate populations four and five. The data generated was then used to estimate parameters of a population model that describes the dynamics of interacting species (cf. below).

**Empirical estimation of population growth rate in presence of interspecific competitors**

To test how evolution in presence or absence of cadmium shaped the growth of populations when competing with other species, we performed a growth rate experiment combining individuals from the no cadmium/cadmium selection regimes on plants with or without cadmium, creating a full orthogonal design: *T. urticae* cadmium / no-cadmium * *T. evansi* cadmium / no-cadmium in plants with / without cadmium, leading to eight experimental conditions. After two generations we quantified the number of adult females produced on each box. In each experimental box we placed two leaves from plants with or without cadmium (box size: 16.5 x 16.5 x 12 cm). Each box was covered with a lid with a gauze-covered opening to allow air circulation. The petiole of the leaves was in a small pot with water, replenished each week. After one mite generation (two weeks), another two leaves were added to the box. By the end of the second generation, we counted the number of adult females from each species in each box. The experiment was done in five blocks, each including one replicate population of each selection regime. For each experimental condition and experimental evolution replicate (i.e. populations one to five) we had 10 boxes (replicates).

**Quantifying the impact of the cadmium environment in populations evolving in the no cadmium selection regime**

To test the impact of the cadmium environment on the intrinsic growth rate of *T. urticae* and *T. evansi* populations that were naïve to this selection pressure, we applied, for each species, a linear model using as dependent variables the estimates of the intrinsic growth rate (estimated from the cxr, see above) of the five experimental replicates of the no-cadmium selection regimes and environment as fixed factor (with two levels, cadmium and no-cadmium).

To verify that plants with cadmium represent a stress for these populations and thus pose a significant selection pressure, we first compared the intrinsic growth rate of naïve populations (evolving on plants without cadmium) on plants with cadmium (Fig S7). Populations of both mite species from the 'no-cadmium' selection regime had significantly lower growth rates in the cadmium than in the no-cadmium environment (*Tetranychus urticae*: $\chi^2_{1,7}$= 116.73, P-value<0.001, *T. evansi*: $\chi^2_{1,7}$=25.149, P-value<0.001, Table S1), confirming our hypothesis.

**Estimating parameters for the Riker model with the cxr package**

To predict how evolution in cadmium affected the probability of coexistence between the two spider-mite species, we used the data from the competitive ability experiment to parameterize the Riker model (1954) that describes the number of individuals produced by two competing species.

In the initial step, we estimated the intrinsic growth rate as well as intraspecific and interspecific competitive abilities using the cxr package (García-Callejas et al., 2022), using three estimation approaches: global estimation, fixed lambda, and the nested estimation method. In addition, we applied the nested method previously used by Matias et al. (2018) with and without the fixed lambda (i.e. lambda estimated directly from the data and inputted in the model to estimate the alphas). Using the parameter estimates obtained from each method, we compared the Euclidean distance between observed and predicted data (see code details in the Git repository). Among the approaches tested, the cxr-based estimates consistently resulted in the smallest average distance between observed and predicted values (Fig. S2), and showed the lowest variation in parameter estimation. Consequently, this method was used in all subsequent analyses.

The cxr package uses optim to estimate model parameters. However, such optimizers need several starting points, primarily because local optimization methods can converge to suboptimal local minima rather than the global minimum. Thus, using multiple starting points allows the optimizer to explore different regions of the solution space, increasing the chances of finding the best possible solution. We detected a problem of suboptimal fitting because of initial conditions when performing the data fitting with the cxr package, in particular when using the pooled data. To mitigate this, we performed a grid search over a defined range of starting values: intrinsic growth rates varied from 0.001 to 2, while intra- and interspecific competitive coefficients ranged from -1 to 1, both using an interval of 0.1.

We used the cxr multifit function to estimate parameters for each combination of starting values for the dataset with each individual replicates and pooled replicates, and for both the cadmium and no-cadmium environments. For each environment and combination of initial values we computed the mean likelihood across the five replicates. For the final analyses we used the starting conditions that showed the best mean likelihood (i.e. minimum) across the five replicates. For the pooled replicates, the initial conditions were chosen based on the initial conditions that showed the best mean likelihood for all selection regimes. The starting points used were: $\lambda_i$=0.9, $\alpha_{ii}$=-0.1, $\alpha_{ij}$=0 for the individual data in the normal and cadmium environment, $\lambda_i$=1.6, $\alpha_{ii}$=0, $\alpha_{ij}$=-0.1 for the pooled data in the normal and cadmium environment. Details of this procedure and the code used are available in the online repository (https://figshare.com/s/f001d9f699a4027d7b62)

### Supplementary Figures

A)

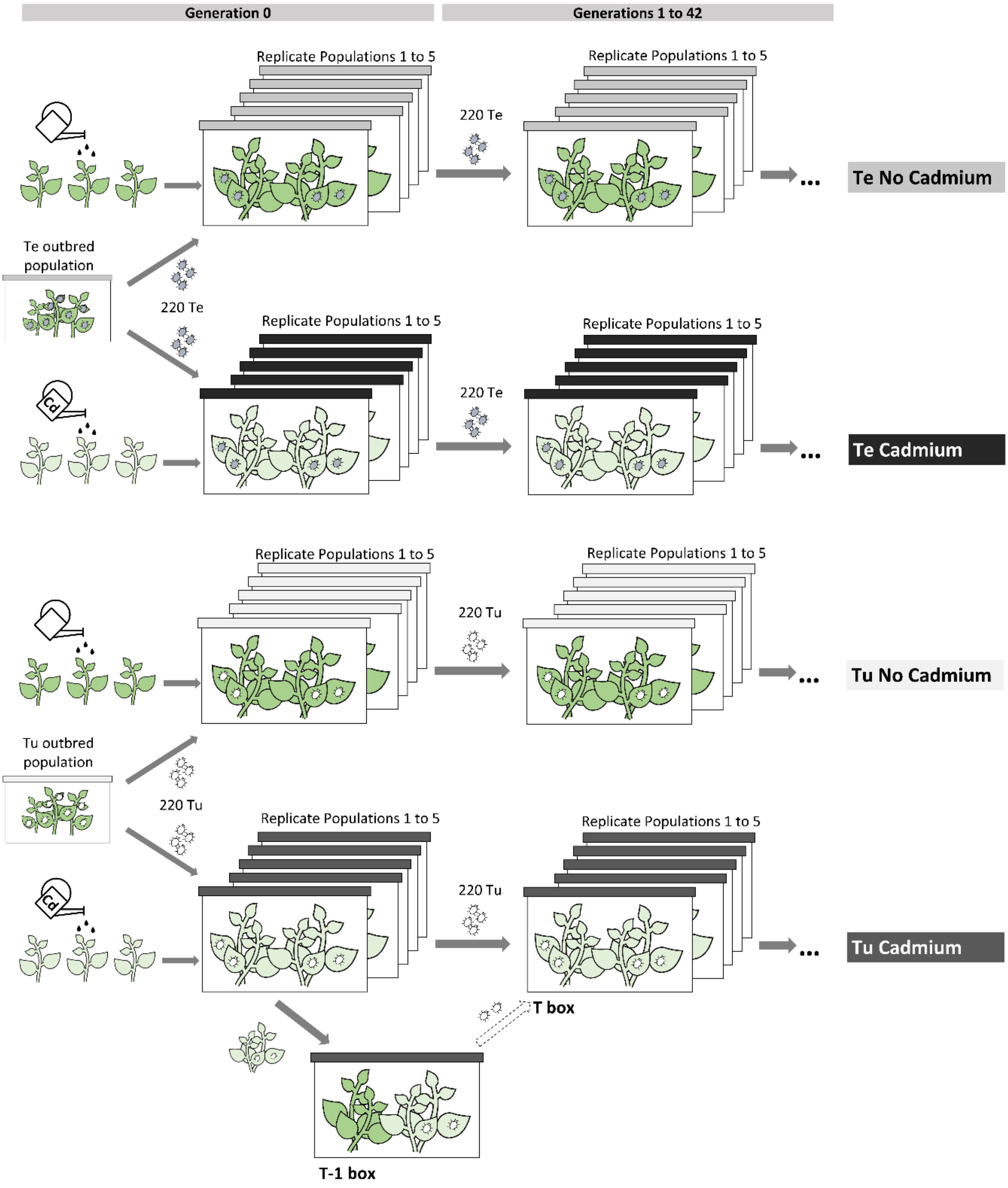

B)

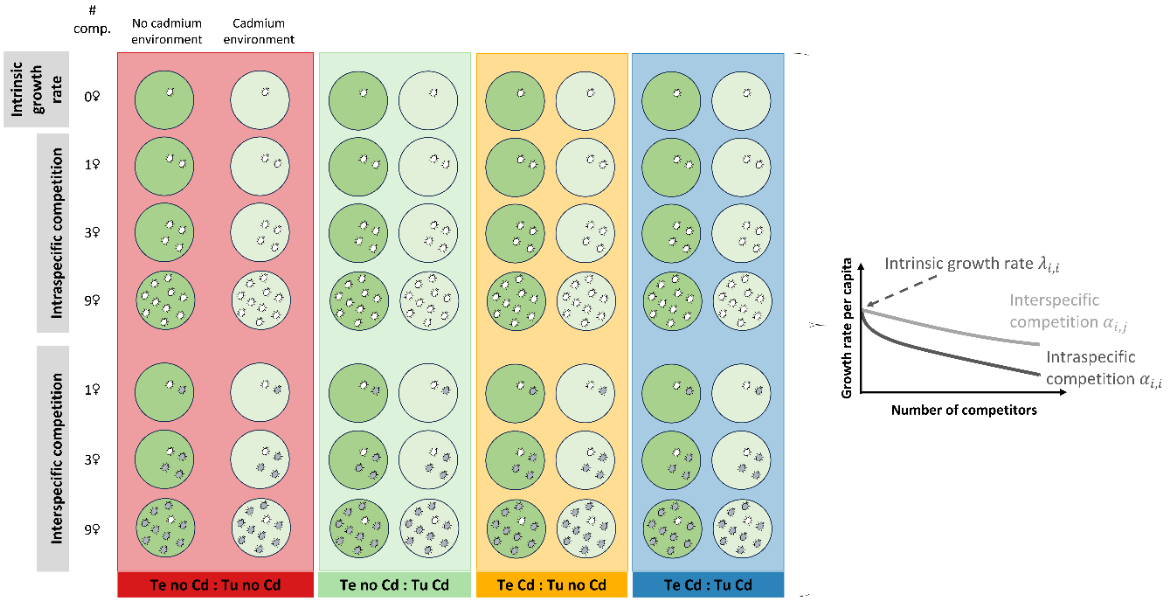

C)

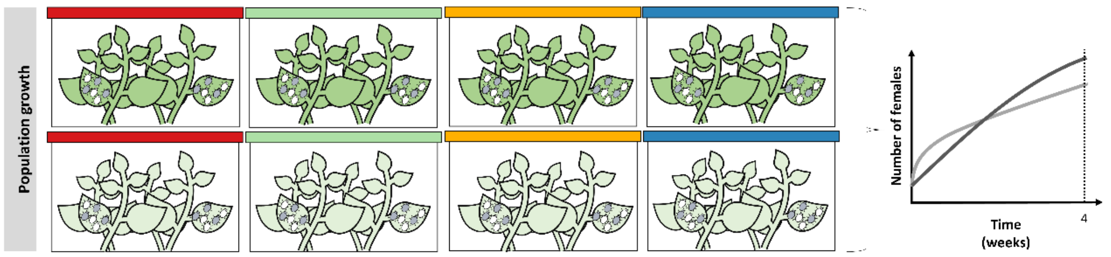

**Figure S1**

**Overview of the experimental procedure. A) Experimental evolution:** 220 females from the T. urticae (Tu) or T. evansi (Te) outbred populations were transferred to create four experimental regimes: No-cadmium (plants grown in soil without cadmium, dark green) and Cadmium (plants grown in soil with 2mM cadmium, light green), for each mite species evolved in absence of interspecific competition. Each selection regime was replicated five times. Every two weeks (roughly corresponding to one spider mite generation), 220 females were transferred from each population to a new box with the same plant treatment. This procedure was followed for 42 mite generations. **B) Experiment to estimate strength of competition:** Females from the four experimental regimes were placed on a leaf disk of a plant grown with (light green) or without (dark green) cadmium. Focal females were exposed to a gradient of intraspecific or interspecific female competitors stemming from the cadmium or no-cadmium selection regimes. In total, four possible combinations of cadmium and no-cadmium selection regimes were performed: Te no-cadmium: Tu no-cadmium (red), Te no-cadmium: Tu cadmium (green), Te cadmium: Tu no-cadmium (yellow), Te cadmium: Tu cadmium (blue). The number of adult female offspring was measured after two weeks to calculate the per capita offspring production (total number of offspring divided by the number of focal females initially added to the patch). These data were then used to parameterize a Ricker model to estimate the intrinsic growth rate and the intra and interspecific competition coefficients. The parameters were then used to estimate 1) the relative impact of intra and interspecific competition by predicting the number of offspring produced under different scenarios (cf. Figure 1 in main text) and 2) the long-term coexistence outcomes of competition between the different selection regimes (cf. Figure 2 in main text). **C) Population experiment:** Six females from each experimental evolution selection regime were placed in a box with two leaves from plants grown with or without cadmium. Boxes were created for the four possible combinations of cadmium and no-cadmium selection regimes: Te no-cadmium: Tu no-cadmium (red), Te no-cadmium: Tu cadmium (green), Te cadmium: Tu no-cadmium (yellow), Te cadmium: Tu cadmium (blue). After two weeks, two more plants were added, and the number of adult female offspring of each mite species was counted four weeks later. Figure adapted from Godinho et al (2024).

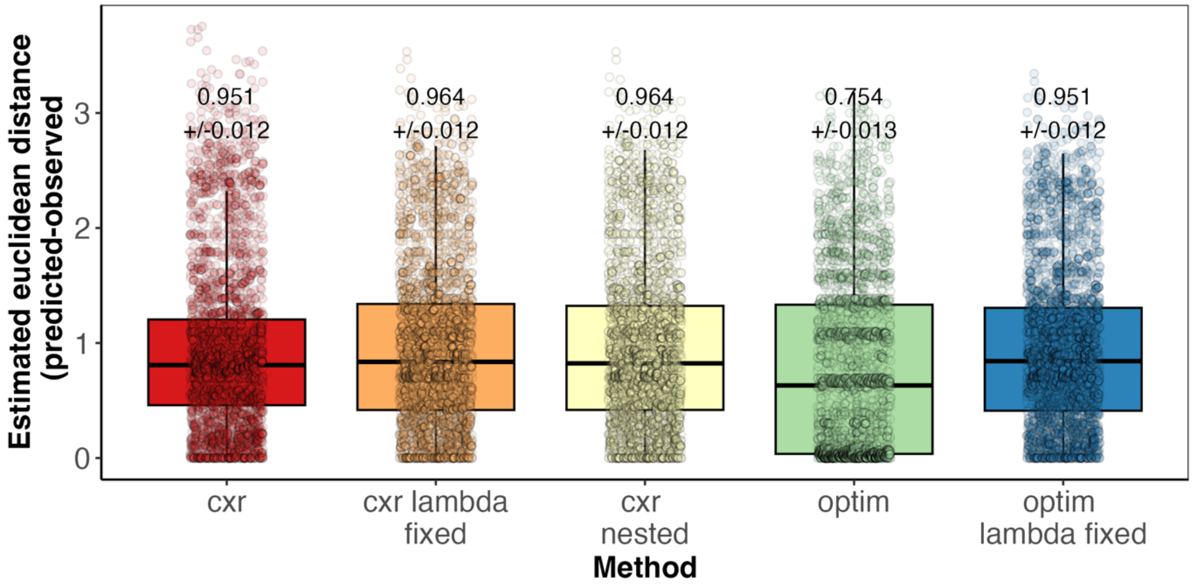

**Figure S2**
**Euclidean distance between predicted and observed estimates for five different fitting approaches (x axis).** Boxplot limits represent first and third quartiles and the whiskers mark points within 1.5*interquartile range. The different methods tested were <u>cxr:</u> estimates fitted using the cxr package; <u>cxr lambda fixed:</u> using the cxr package with the intrinsic growth rate estimates directly obtained from the single female assays; <u>cxr nested:</u> using the cxr package but estimating first intrinsic growth rate, then intraspecific competitive ability and then the interspecific competitive ability; <u>optim:</u> using the method described in Matias et al (2018); <u>optim lambda fixed:</u> using the same method but with the intrinsic growth rate estimates directly obtained from the single female assays. Numbers in the plot indicate average distance for each method and their standard error. The optim method shows on average the smallest distance between predicted and observed, however it also shows some of the highest variation and error associated with parameter estimation, so we used the cxr package that shows the second lowest average distance. Details for the methods are available in the git repository.

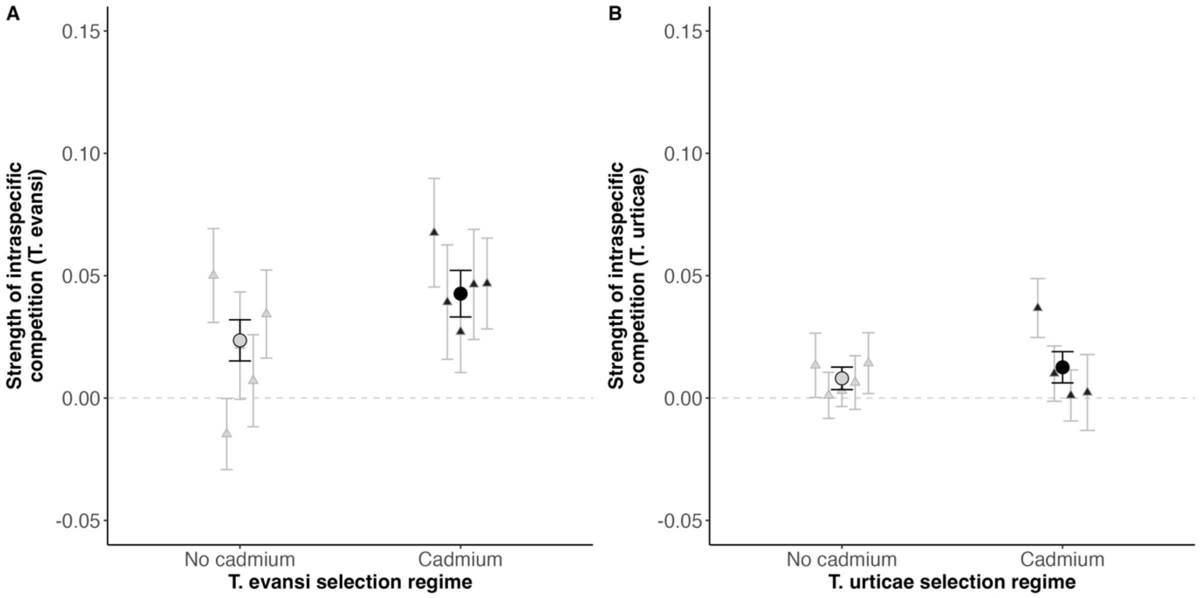

**Figure S3**

**Sensitivity to intraspecific competition for T. evansi and T. urticae selection regimes in the cadmium environment.** The y-axis corresponds to the strength of intraspecific competition experienced by *T. evansi* (A) or *T. urticae* (B). No cadmium and cadmium selection regimes are represented in light and dark colours, respectively. Error bars were calculated based on standard error obtained from 1000 bootstrap samples. Circles correspond to the parameters estimated from all replicates pooled and triangles to parameters estimated from each replicate. Note that the scales are different between the two panels.

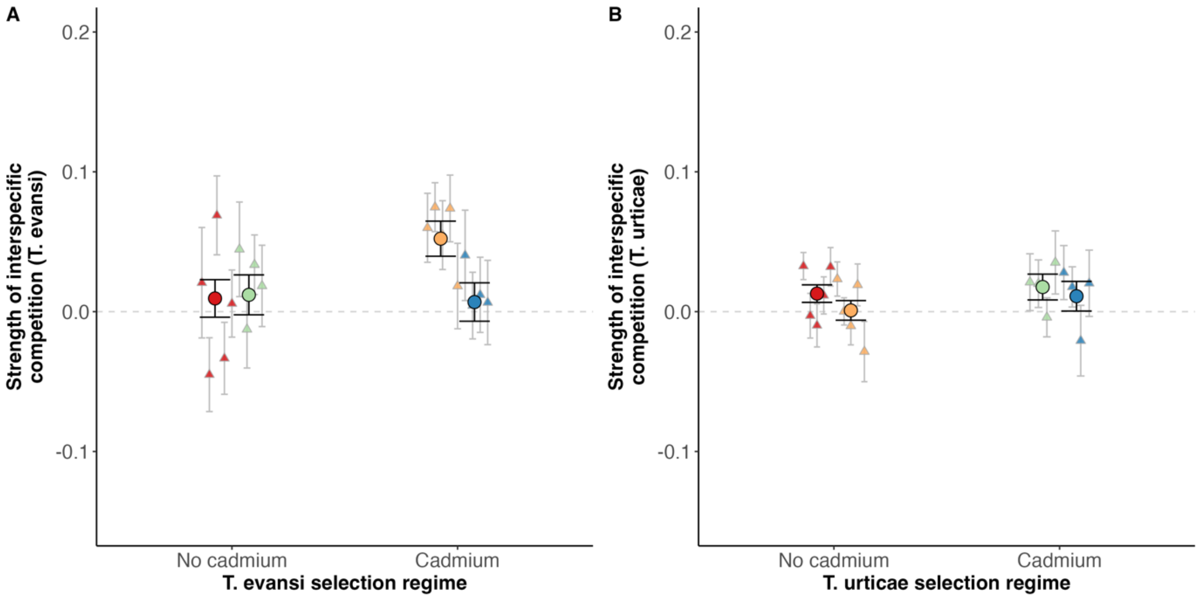

**Figure S4**

**Sensitivity to interspecific competition for *T. evansi* and *T. urticae* selection regimes in the cadmium environment.** The y-axis corresponds to the strength of interspecific competition experienced by *T. evansi* (A) or *T. urticae* (B). Colors indicate the four possible combinations of cadmium and no-cadmium selection regimes were performed: Te no cadmium: Tu no cadmium (red), Te no cadmium: Tu cadmium (green), Te cadmium: Tu no cadmium (yellow), Te cadmium: Tu cadmium (blue). Error bars were calculated based on standard error obtained from 1000 bootstrap samples. Circles correspond to the parameters estimated from all replicates pooled and triangles to parameters estimated from each replicate. Note that the scales are different between the two panels.

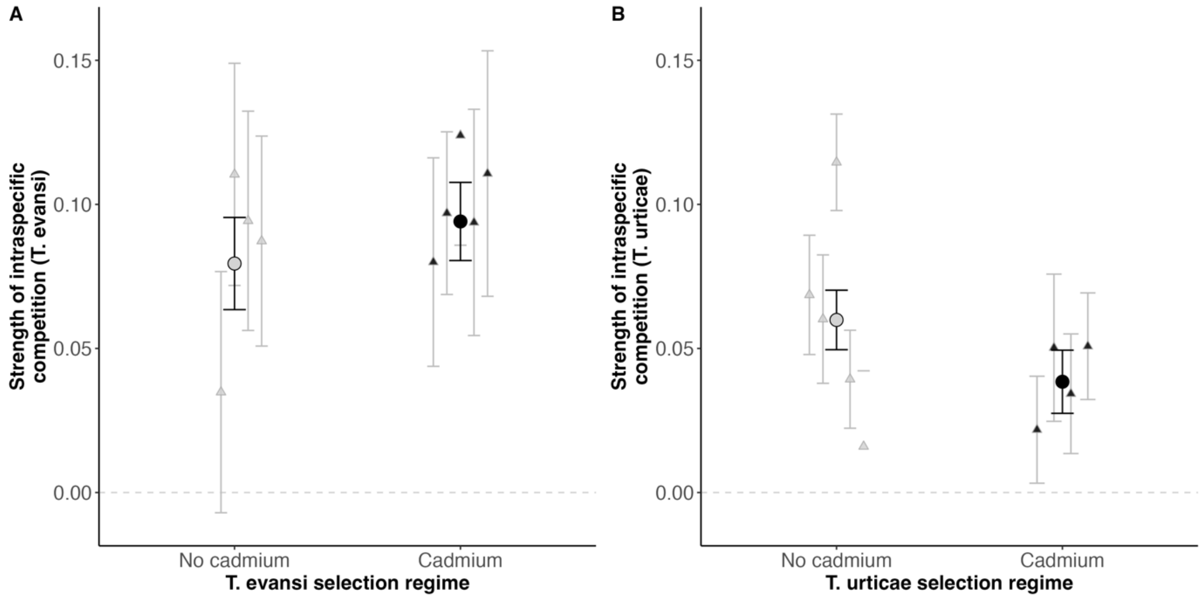

**Figure S5**

**Sensitivity to intraspecific competition for *T. evansi* and *T. urticae* selection regimes in the no cadmium environment.** The y-axis corresponds to the strength of intraspecific competition experienced by *T. evansi* (A) or *T. urticae* (B). No-cadmium and cadmium selection regimes are represented in light and dark colours, respectively. Error bars were calculated based on standard error obtained from 1000 bootstrap samples. Circles correspond to the parameters estimated from all replicates pooled and triangles to parameters estimated from each replicate. Note that the scales are different between the two panels.

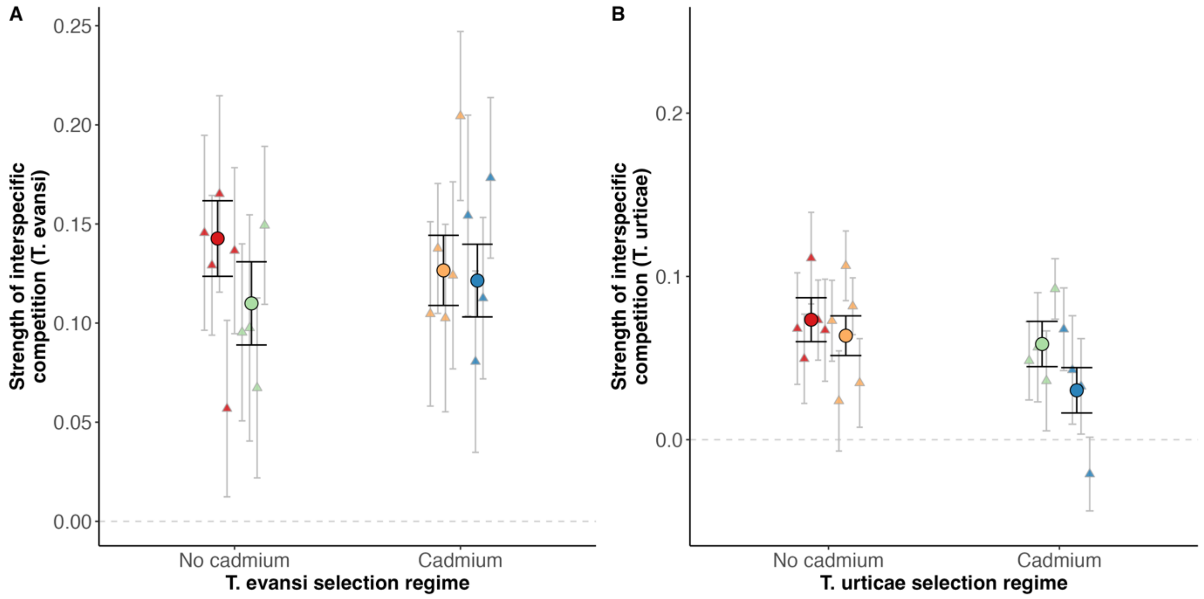

**Figure S6**

**Sensitivity to interspecific competition for *T. evansi* and *T. urticae* selection regimes in the no cadmium environment.** The y-axis corresponds to the strength of interspecific competition experienced by *T. evansi* (A) or *T. urticae* (B). Colors indicate the four possible combinations of cadmium and no-cadmium selection regimes were performed: Te no-cadmium: Tu no-cadmium (red), Te no-cadmium: Tu cadmium (green), Te cadmium: Tu no-cadmium (yellow), Te cadmium: Tu cadmium (blue). Error bars were calculated based on standard error obtained from 1000 bootstrap samples. Circles correspond to the parameters estimated from all replicates pooled and triangles to parameters estimated from each replicate. Note that the scales are different between the two panels.

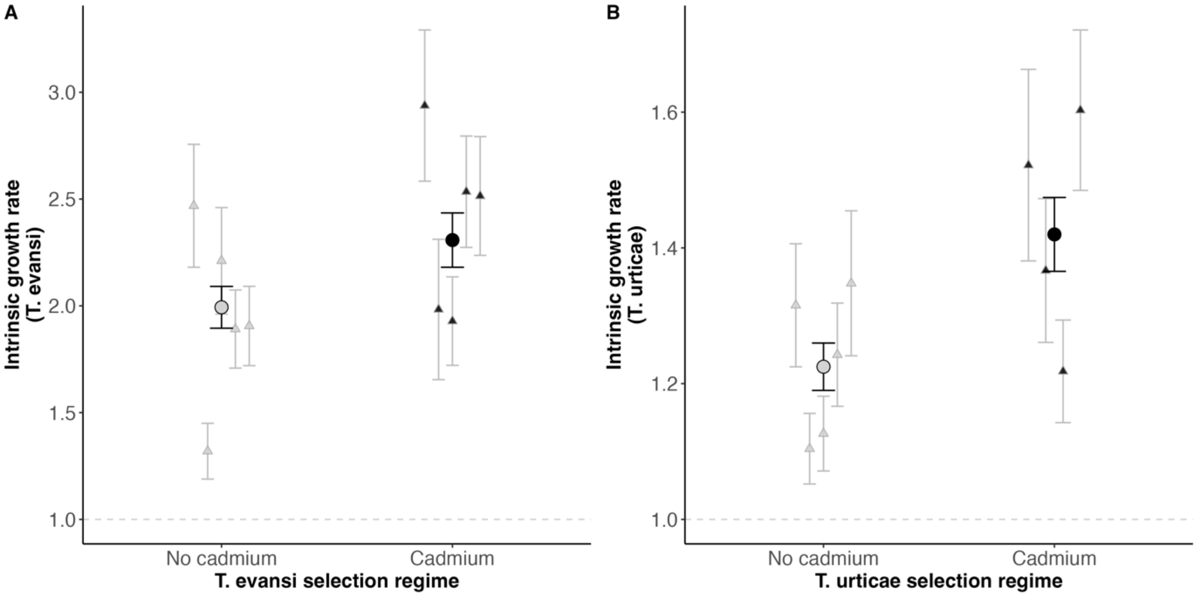

**Figure S7**
**Intrinsic growth rate of *T. evansi* (A) and *T. urticae* (B) from the cadmium and the no cadmium selection regimes, when tested in the cadmium environment.** No-cadmium and cadmium selection regimes are represented in light and dark colours, respectively. Error bars were calculated based on standard error obtained from 1000 bootstrap samples. Circles correspond to the parameters estimated from all replicates pooled and triangles to parameters estimated from each replicate. Note that the scales are different between the two panels.

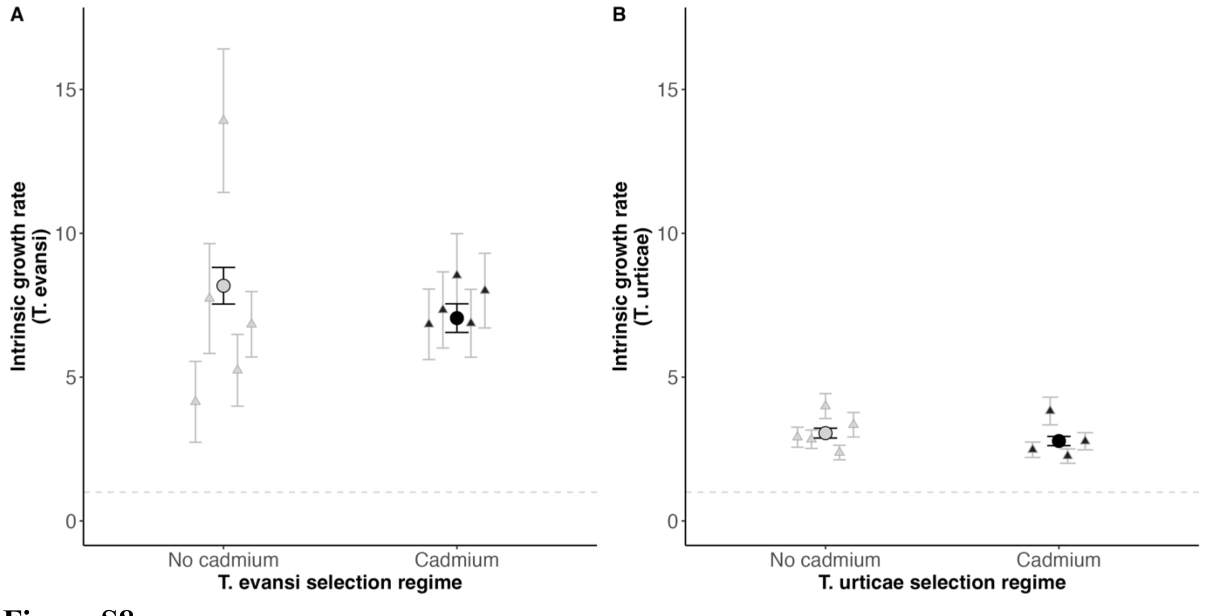

**Figure S8**
**– Intrinsic growth rate for *T. evansi* (A) and *T. urticae* (B) selection regimes in the no cadmium environment.** No-cadmium and cadmium selection regimes are represented in light

and dark colours, respectively. Error bars were calculated based on standard error obtained from 1000 bootstrap samples. Circles correspond to the parameters estimated from all replicates pooled and triangles to parameters estimated from each replicate. Note that the scales are different between the two panels.

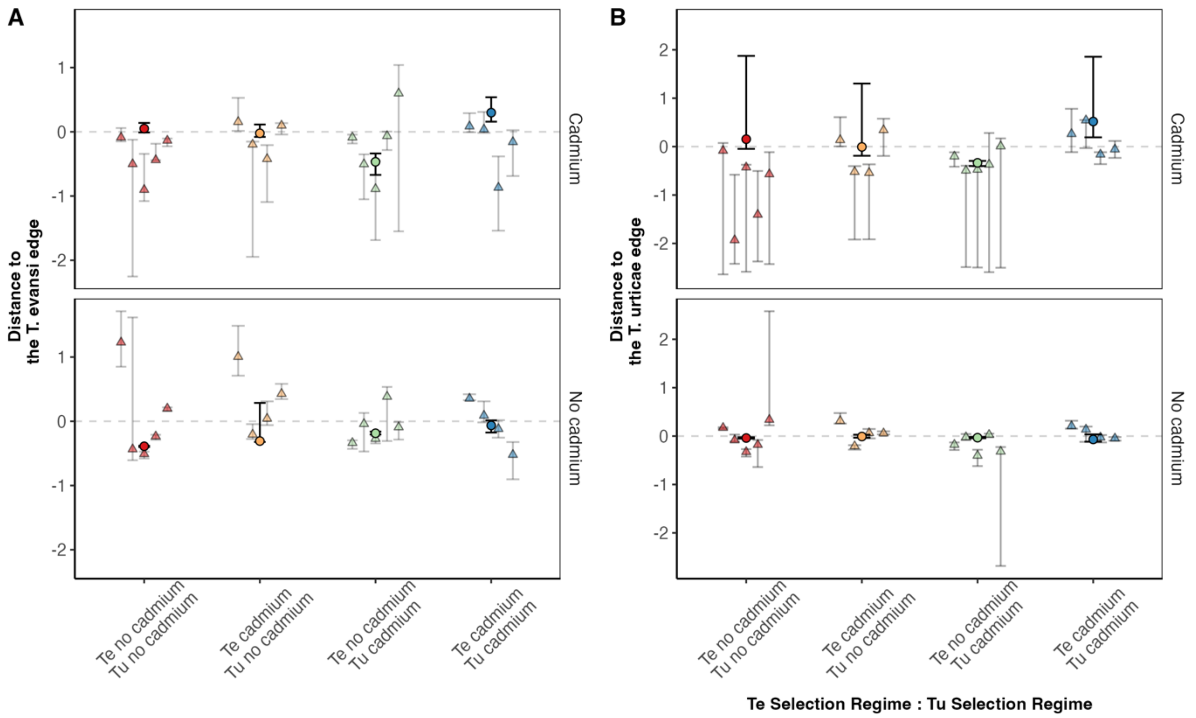

**Figure S9**

**– Distance to the two edges of the feasibility domain for the different combinations of cadmium and no-cadmium selection regimes of *T. urticae* (Tu) and *T. evansi* (Te) in the cadmium and the no-cadmium environment.** Distance between the realized growth rates and the edges of the feasibility domain for *T. evansi* (A) or *T. urticae* (B, in the cadmium (upper panels) and no-cadmium (lower panels) environments for the different treatments (i.e. combinations of no cadmium or cadmium selection regimes, cf. colour codes). Positive distances indicate that the vector of growth rates is inside of the feasibility domain (i.e., coexistence is possible), and negative distances indicate that the vector is outside of the feasibility domain (one species is excluded). Circles correspond to the distance calculated with data from all replicates pooled and triangles to distance calculated per replicate. Note the differences in scales between panels. Confidence intervals describe the difference between the vector of intrinsic growth rates and the edges of the largest or smallest feasibility domain, which were obtained from the lower and upper estimates of the parameters as described in Figure 2. The pooled data estimates were obtained from data from all experimental evolution replicates to increase statistical power and to account for the variation between replicates.

#### Supplementary Tables

**Table S1**

Estimated intrinsic growth rate (λ), intra- (α intra) and interspecific (α inter) competitive ability using data from the pooled replicates (A) or each replicate separately (B). Confidence intervals for each estimate were obtained using 1000 bootstrap samples (from cxr package).

A) Pooled replicates

| Selection Regime T. urticae | Selection Regime T. evansi | Env | λ T. urticae (Lower : Upper CI) | λ T. evansi (Lower : Upper CI) | α intra T. urticae (Lower - Upper CI) | α intra T. evansi (Lower : Upper CI) | α inter T. urticae (Lower : UpperCI) | α inter T. evansi (Lower : Upper CI) |
|---|---|---|---|---|---|---|---|---|
| No cadmium | No cadmium | No cadmium | 3.053 (2.882 - 3.224) | 8.178 (7.541 - 8.814) | 0.06 (0.05 - 0.07) | 0.079 (0.063 - 0.095) | 0.074 (0.06 - 0.087) | 0.143 (0.124 - 0.162) |
| Cadmium | No cadmium | No cadmium | 2.779 (2.618 - 2.94) | 8.178 (7.541 - 8.814) | 0.038 (0.027 - 0.049) | 0.079 (0.063 - 0.095) | 0.059 (0.045 - 0.072) | 0.11 (0.089 - 0.131) |
| No cadmium | Cadmium | No cadmium | 3.053 (2.882 - 3.224) | 7.052 (6.555 - 7.549) | 0.06 (0.05 - 0.07) | 0.094 (0.081 - 0.108) | 0.064 (0.052 - 0.076) | 0.127 (0.109 - 0.144) |
| Cadmium | Cadmium | No cadmium | 2.779 (2.618 - 2.94) | 7.052 (6.555 - 7.549) | 0.038 (0.027 - 0.049) | 0.094 (0.081 - 0.108) | 0.03 (0.016 - 0.044) | 0.121 (0.103 - 0.14) |
| No cadmium | No cadmium | Cadmium | 1.225 (1.19 - 1.26) | 1.993 (1.895 - 2.091) | 0.008 (0.003 - 0.013) | 0.024 (0.015 - 0.032) | 0.013 (0.007 - 0.019) | 0.009 (-0.004 - 0.023) |
| Cadmium | No cadmium | Cadmium | 1.42 (1.365 - 1.474) | 1.993 (1.895 - 2.091) | 0.013 (0.006 - 0.019) | 0.024 (0.015 - 0.032) | 0.018 (0.008 - 0.027) | 0.012 (-0.002 - 0.026) |
| No cadmium | Cadmium | Cadmium | 1.225 (1.19 - 1.26) | 2.308 (2.181 - 2.435) | 0.008 (0.003 - 0.013) | 0.043 (0.033 - 0.052) | 0.001 (-0.006 - 0.008) | 0.052 (0.04 - 0.065) |
| Cadmium | Cadmium | Cadmium | 1.42 (1.365 - 1.474) | 2.308 (2.181 - 2.435) | 0.013 (0.006 - 0.019) | 0.043 (0.033 - 0.052) | 0.011 (0 - 0.022) | 0.007 (-0.007 - 0.021) |

B) Each replicate

| **Selection *T. urticae*** | **Selection *T. evansi*** | **Environment** | **Replicate** | **ƛ T. urticae (Lower - Upper bounds)** | **ƛ T. evansi (Lower - Upper bounds)** | **α intra T. urticae (Lower - Upper bounds)** | **α intra T. evansi (Lower - Upper bounds)** | **α inter T. urticae (Lower - Upper bounds)** | **α inter T. evansi (Lower - Upper bounds)** |
|---|---|---|---|---|---|---|---|---|---|
| No cadmium | No cadmium | No cadmium | 1 | 2.911 (2.562 - 3.26) | 4.142 (2.737 - 5.548) | 0.069 (0.048 - 0.089) | -0.096 (-0.14 - -0.052) | 0.068 (0.034 - 0.102) | 0.146 (0.096 - 0.195) |
| Cadmium | No cadmium | No cadmium | 1 | 2.476 (2.209 - 2.744) | 4.142 (2.737 - 5.548) | 0.022 (0.003 - 0.04) | -0.096 (-0.14 - -0.052) | 0.048 (0.024 - 0.072) | 0.095 (0.051 - 0.14) |
| No cadmium | Cadmium | No cadmium | 1 | 2.911 (2.562 - 3.26) | 6.836 (5.612 - 8.061) | 0.069 (0.048 - 0.089) | 0.08 (0.044 - 0.116) | 0.073 (0.048 - 0.098) | 0.105 (0.058 - 0.151) |
| Cadmium | Cadmium | No cadmium | 1 | 2.476 (2.209 - 2.744) | 6.836 (5.612 - 8.061) | 0.022 (0.003 - 0.04) | 0.08 (0.044 - 0.116) | 0.068 (0.042 - 0.093) | 0.154 (0.104 - 0.205) |
| No cadmium | No cadmium | No cadmium | 2 | 2.841 (2.521 - 3.16) | 7.735 (5.826 - 9.643) | 0.06 (0.038 - 0.082) | 0.035 (-0.007 - 0.077) | 0.05 (0.022 - 0.077) | 0.129 (0.094 - 0.164) |
| No cadmium | Cadmium | No cadmium | 2 | 2.841 (2.521 - 3.16) | 7.336 (6.011 - 8.66) | 0.06 (0.038 - 0.082) | 0.097 (0.069 - 0.125) | 0.024 (-0.007 - 0.054) | 0.138 (0.105 - 0.17) |
| No cadmium | No cadmium | No cadmium | 3 | 3.994 (3.559 - 4.429) | 13.915 (11.421 - 16.409) | 0.115 (0.098 - 0.131) | 0.11 (0.072 - 0.149) | 0.111 (0.083 - 0.139) | 0.165 (0.116 - 0.215) |
| Cadmium | No cadmium | No cadmium | 3 | 3.822 (3.34 - 4.303) | 13.915 (11.421 - 16.409) | 0.05 (0.025 - 0.076) | 0.11 (0.072 - 0.149) | 0.057 (0.023 - 0.09) | 0.098 (0.04 - 0.155) |
| No cadmium | Cadmium | No cadmium | 3 | 3.994 (3.559 - 4.429) | 8.535 (7.083 - 9.988) | 0.115 (0.098 - 0.131) | 0.124 (0.086 - 0.162) | 0.106 (0.085 - 0.128) | 0.103 (0.055 - 0.15) |
| Cadmium | Cadmium | No cadmium | 3 | 3.822 (3.34 - 4.303) | 8.535 (7.083 - 9.988) | 0.05 (0.025 - 0.076) | 0.124 (0.086 - 0.162) | 0.043 (0.01 - 0.076) | 0.081 (0.035 - 0.126) |
| No cadmium | No cadmium | No cadmium | 4 | 2.378 (2.125 - 2.632) | 5.239 (3.991 - 6.486) | 0.039 (0.022 - 0.056) | 0.094 (0.056 - 0.132) | 0.073 (0.049 - 0.098) | 0.057 (0.012 - 0.101) |
| Cadmium | No cadmium | No cadmium | 4 | 2.257 (2.008 - 2.506) | 5.239 (3.991 - 6.486) | 0.034 (0.014 - 0.055) | 0.094 (0.056 - 0.132) | 0.036 (0.005 - 0.067) | 0.067 (0.022 - 0.113) |
| No cadmium | Cadmium | No cadmium | 4 | 2.378 (2.125 - 2.632) | 6.872 (5.692 - 8.051) | 0.039 (0.022 - 0.056) | 0.094 (0.054 - 0.133) | 0.082 (0.064 - 0.099) | 0.124 (0.077 - 0.171) |
| Cadmium | Cadmium | No cadmium | 4 | 2.257 (2.008 - 2.506) | 6.872 (5.692 - 8.051) | 0.034 (0.014 - 0.055) | 0.094 (0.054 - 0.133) | 0.033 (0.003 - 0.062) | 0.113 (0.072 - 0.153) |
| No cadmium | No cadmium | No cadmium | 5 | 3.345 (2.919 - 3.771) | 6.837 (5.698 - 7.975) | 0.016 (-0.01 - 0.042) | 0.087 (0.051 - 0.124) | 0.067 (0.036 - 0.098) | 0.137 (0.095 - 0.178) |
| Cadmium | No cadmium | No cadmium | 5 | 2.771 (2.473 - 3.069) | 6.837 (5.698 - 7.975) | 0.051 (0.032 - 0.069) | 0.087 (0.051 - 0.124) | 0.092 (0.074 - 0.111) | 0.149 (0.109 - 0.189) |
| No cadmium | Cadmium | No cadmium | 5 | 3.345 (2.919 - 3.771) | 8.007 (6.71 - 9.304) | 0.016 (-0.01 - 0.042) | 0.111 (0.068 - 0.153) | 0.035 (0.008 - 0.062) | 0.204 (0.162 - 0.247) |

| | | | | | | | | | |
|---|---|---|---|---|---|---|---|---|---|
| Cadmium | Cadmium | No cadmium | 5 | 2.771 (2.473 - 3.069) | 8.007 (6.71 - 9.304) | 0.051 (0.032 - 0.069) | 0.111 (0.068 - 0.153) | -0.021 (-0.044 - 0.002) | 0.173 (0.133 - 0.214) |
| No cadmium | No cadmium | Cadmium | 1 | 1.315 (1.225 - 1.406) | 2.468 (2.181 - 2.756) | 0.013 (0 - 0.027) | 0.05 (0.031 - 0.069) | 0.033 (0.023 - 0.042) | 0.021 (-0.019 - 0.06) |
| Cadmium | No cadmium | Cadmium | 1 | 1.522 (1.381 - 1.663) | 2.468 (2.181 - 2.756) | 0.037 (0.025 - 0.049) | 0.05 (0.031 - 0.069) | 0.021 (0.001 - 0.041) | 0.045 (0.011 - 0.078) |
| No cadmium | Cadmium | Cadmium | 1 | 1.315 (1.225 - 1.406) | 2.938 (2.584 - 3.292) | 0.013 (0 - 0.027) | 0.068 (0.045 - 0.09) | 0.023 (0.011 - 0.036) | 0.06 (0.035 - 0.085) |
| Cadmium | Cadmium | Cadmium | 1 | 1.522 (1.381 - 1.663) | 2.938 (2.584 - 3.292) | 0.037 (0.025 - 0.049) | 0.068 (0.045 - 0.09) | 0.028 (0.009 - 0.047) | 0.04 (0.008 - 0.073) |
| No cadmium | No cadmium | Cadmium | 2 | 1.104 (1.052 - 1.156) | 1.319 (1.189 - 1.45) | 0.001 (-0.008 - 0.011) | -0.015 (-0.029 - 0) | -0.003 (-0.019 - 0.013) | -0.045 (-0.071 - - 0.019) |
| No cadmium | Cadmium | Cadmium | 2 | 1.104 (1.052 - 1.156) | 1.983 (1.655 - 2.312) | 0.001 (-0.008 - 0.011) | 0.039 (0.016 - 0.063) | 0 (-0.01 - 0.01) | 0.075 (0.057 - 0.092) |
| No cadmium | No cadmium | Cadmium | 3 | 1.126 (1.071 - 1.182) | 2.211 (1.962 - 2.46) | 0.003 (-0.003 - 0.009) | 0.021 (-0.001 - 0.043) | -0.01 (-0.025 - 0.006) | 0.069 (0.041 - 0.097) |
| Cadmium | No cadmium | Cadmium | 3 | 1.367 (1.261 - 1.473) | 2.211 (1.962 - 2.46) | 0.01 (-0.001 - 0.021) | 0.021 (-0.001 - 0.043) | 0.02 (0.003 - 0.037) | -0.013 (-0.04 - 0.015) |
| No cadmium | Cadmium | Cadmium | 3 | 1.126 (1.071 - 1.182) | 1.929 (1.721 - 2.136) | 0.003 (-0.003 - 0.009) | 0.027 (0.01 - 0.044) | -0.01 (-0.024 - 0.003) | 0.055 (0.03 - 0.079) |
| Cadmium | Cadmium | Cadmium | 3 | 1.367 (1.261 - 1.473) | 1.929 (1.721 - 2.136) | 0.01 (-0.001 - 0.021) | 0.027 (0.01 - 0.044) | 0.018 (0.003 - 0.032) | 0.004 (-0.019 - 0.028) |
| No cadmium | No cadmium | Cadmium | 4 | 1.243 (1.167 - 1.319) | 1.891 (1.708 - 2.074) | 0.006 (-0.005 - 0.017) | 0.007 (-0.012 - 0.026) | 0.012 (-0.002 - 0.025) | -0.033 (-0.059 - - 0.008) |
| Cadmium | No cadmium | Cadmium | 4 | 1.218 (1.143 - 1.294) | 1.891 (1.708 - 2.074) | 0.001 (-0.009 - 0.011) | 0.007 (-0.012 - 0.026) | -0.004 (-0.018 - 0.01) | 0.034 (0.012 - 0.055) |
| No cadmium | Cadmium | Cadmium | 4 | 1.243 (1.167 - 1.319) | 2.534 (2.274 - 2.795) | 0.006 (-0.005 - 0.017) | 0.046 (0.024 - 0.069) | 0.019 (0.004 - 0.034) | 0.074 (0.05 - 0.098) |
| Cadmium | Cadmium | Cadmium | 4 | 1.218 (1.143 - 1.294) | 2.534 (2.274 - 2.795) | 0.001 (-0.009 - 0.011) | 0.046 (0.024 - 0.069) | -0.021 (-0.046 - 0.005) | 0.012 (-0.015 - 0.039) |
| No cadmium | No cadmium | Cadmium | 5 | 1.348 (1.241 - 1.455) | 1.906 (1.72 - 2.091) | 0.014 (0.002 - 0.027) | 0.034 (0.016 - 0.052) | 0.032 (0.018 - 0.046) | 0.006 (-0.018 - 0.03) |
| Cadmium | No cadmium | Cadmium | 5 | 1.603 (1.485 - 1.721) | 1.906 (1.72 - 2.091) | 0.002 (-0.013 - 0.018) | 0.034 (0.016 - 0.052) | 0.035 (0.013 - 0.058) | 0.018 (-0.011 - 0.047) |
| No cadmium | Cadmium | Cadmium | 5 | 1.348 (1.241 - 1.455) | 2.514 (2.236 - 2.793) | 0.014 (0.002 - 0.027) | 0.047 (0.028 - 0.065) | -0.029 (-0.05 - - 0.007) | 0.018 (-0.012 - 0.049) |
| Cadmium | Cadmium | Cadmium | 5 | 1.603 (1.485 - 1.721) | 2.514 (2.236 - 2.793) | 0.002 (-0.013 - 0.018) | 0.047 (0.028 - 0.065) | 0.02 (-0.003 - 0.044) | 0.007 (-0.024 - 0.037) |

**Table S2**

**Bootstrap analyses** to test for differences in intrinsic growth rate (Lambda), intraspecific and interspecific competition 1) between no-cadmium and cadmium environments (only in the evolved without cadmium regimes); 2) between evolved with cadmium and evolved without cadmium regimes on the cadmium environment and 3) between evolved with cadmium and evolved without cadmium regimes on the no cadmium environment. The models were applied separately to each species (*T. urticae* and *T. evansi*). For each model we computed the number of times that a P-value lower or equal than 0.05 was obtained. Bootstrap was done using 10000 samples with replacement.

| Test | Species | Selection regimes | Environment | Term | Observed P-value | Probability of obtaining a P-value <=0.05 |
|---|---|---|---|---|---|---|
| 1) What is the impact of cadmium on spider-mite performance for the selection regimes that evolved without cadmium? | *T. urticae* | Evolved without cadmium | Cadmium vs no cadmium | Lambda | < **0.0001** | 0.0013 |
| | | | | Intraspecific alpha | < **0.0001** | 0.0101 |
| | | | | Interspecific alpha | < **0.0001** | 0.0031 |
| | *T. evansi* | | | Lambda | < **0.0001** | 0.0012 |
| | | | | Intraspecific alpha | 0.4516 | 0.5271 |
| | | | | Interspecific alpha | **<0.0001** | 0.0041 |
| 2) What is the impact of evolution with cadmium on spider-mite performance on the cadmium environment? | *T. urticae* | Evolved with cadmium vs Evolved without cadmium | Cadmium | Lambda | **0.0142** | 0.0704 |
| | | | | Intraspecific alpha | 0.4657 | 0.6037 |
| | | | | Interspecific Focal | 0.6512 | 0.4108 |
| | | | | Interspecific Competitor | 0.2839 | 0.3205 |
| | | | | Interspecific Interaction | 0.7563 | 0.7879 |
| | *T. evansi* | | | Lambda | 0.0945 | 0.1626 |
| | | | | Intraspecific alpha | **0.0270** | 0.0823 |
| | | | | Interspecific Focal | **0.0019** | 0.0836 |
| | | | | Interspecific Competitor | 0.3307 | 0.4346 |
| | | | | Interspecific Interaction | **0.0231** | 0.0667 |
| 3) What is the impact of evolution with cadmium on spider-mite performance on the no cadmium environment? | *T. urticae* | Evolved with cadmium vs Evolved without cadmium | No cadmium | Lambda | 0.48 | 0.5454 |
| | | | | Intraspecific alpha | 0.2377 | 0.3413 |
| | | | | Interspecific Focal | 0.3846 | 0.1079 |
| | | | | Interspecific Competitor | 0.5552 | 0.2282 |
| | | | | Interspecific Interaction | 0.4764 | 0.5469 |
| | *T. evansi* | | | Lambda | 0.9688 | 0.9744 |
| | | | | Intraspecific alpha | 0.1107 | 0.1803 |

| | | | | | | |
|---|---|---|---|---|---|---|
| | | | | Interspecific Focal | 0.7195 | 0.3849 |
| | | | | Interspecific Competitor | 0.3055 | 0.4067 |
| | | | | Interspecific Interaction | 0.552 | 0.6151 |

**Table S3**

**Analyses of differences in interspecific competition in the cadmium environment**. (A) Summary of the ANOVA (type III) to estimate the effect of evolving on plants with cadmium on the strength of interspecific competition for *T. urticae* (Tu) and *T. evansi* (Te). (B) Contrasts between the strength of interspecific competition for the *T. evansi* cadmium and non-cadmium selection regimes. Contrasts were obtained using the emmeans function, from the linear model including the selection regimes of focal and competitor individuals as well as their interaction.

A)

| | ***T. urticae*** | | ***T. evansi*** | |
|---|---|---|---|---|
| **Parameter** | **Chisq** | **Pr(>Chisq)** | **Chisq** | **Pr(>Chisq)** |
| Tu Regime | 0.2043 | 0.6512 | 0.9461 | 0.3307 |
| Te Regime | 1.1483 | 0.2839 | 9.6376 | 0.0019** |
| Tu Regime: Te Regime | 0.0963 | 0.7563 | 5.1630 | 0.0231* |

* 0.05 >= P-value > 0.01; ** 0.01 >= P-value > 0.001; *** P-value < 0.001

B)

| **Contrasts *T. evansi*** | **Estimate** | **T ratio** | **P-value** |
|---|---|---|---|
| Te no cadmium: Tu no cadmium - Te cadmium: Tu no cadmium | -0.05293 | -3.104 | 0.0369* |
| Te no cadmium: Tu no cadmium - Te no cadmium: Tu cadmium | -0.01759 | -0.973 | 0.7670 |
| Te no cadmium: Tu no cadmium - Te cadmium: Tu cadmium | -0.01241 | -0.686 | 0.9005 |
| Te cadmium: Tu no cadmium - Te no cadmium: Tu cadmium | 0.03534 | 1.954 | 0.2541 |
| Te cadmium: Tu no cadmium - Te cadmium: Tu cadmium | 0.04052 | 2.241 | 0.1635 |
| Te no cadmium: Tu cadmium - Te cadmium: Tu cadmium | 0.00518 | 0.272 | 0.9926 |

* 0.05 >= P-value > 0.01; ** 0.01 >= P-value > 0.001; *** P-value < 0.001

**Table S4**

**Analyses of differences in interspecific competition in the no-cadmium environment.** Summary of the ANOVA (type III) to estimate the effect of evolving in cadmium on the strength of interspecific competition for *T. urticae* (Tu) and *T. evansi* (Te). The linear model included the selection regimes of focal and competitor individuals as well as their interaction.

| | *T. urticae* | | *T. evansi* | |
|---|---|---|---|---|
| **Parameter** | **Chisq** | **Pr(>Chisq)** | **Chisq** | **Pr(>Chisq)** |
| Tu Regime | 0.7561 | 0.3846 | 1.0499 | 0.3055 |
| Te Regime | 0.3481 | 0.5552 | 0.1290 | 0.7195 |
| Tu Regime: Te Regime | 0.5071 | 0.4764 | 0.3481 | 0.5552 |

**Table S5**
**Analyses of the impact of the cadmium environment on the growth rate, intra and interspecific competition of the no cadmium selection regimes.** Summary of the linear model to estimate the effect of the cadmium environment on the growth rate, strength of intra and interspecific competition for T. urticae (Tu) and T. evansi (Te). The general linear (with the gamma distribution for the growth rate) and linear models (for intra and interspecific competition) included the environment as a fixed factor. Estimate refers to the difference in trait values between the no cadmium and the cadmium environments (i.e. $trait_{No\ cadmium}$- $trait_{Cadmium}$).

| | *T. urticae* | | | *T. evansi* | | |
|---|---|---|---|---|---|---|
| **Trait** | **Estimate** | **Z-value** | **Pr(>\|z\|)** | **Estimate** | **Chisq** | **Pr(>\|z\|)** |
| Growth rate | 0.9245 | 10.804 | < 2e-16 *** | 1.3521 | 6.497 | < 2e-16 *** |
| Intraspecific competition | 0.0521 | 3.494 | 0.00048*** | 0.0265 | 0.753 | 0.452 |
| Interspecific competition | 0.0611 | 5.102 | 3.36e-07 *** | 0.1233 | 5.015 | 5.31e-07 *** |

**Table S6**
**Average proportion of *T. evansi* (Te) females obtained from the experiment to estimate the growth rate of populations with both intra and interspecific competitors.** Each treatment corresponds to a combination of selection regimes (no cadmium Te: no-cadmium Tu, cadmium Te: no-cadmium Tu, no-cadmium Te: cadmium Tu and cadmium Te: cadmium Tu) and was composed of 10 replicate populations. Each box was initialized with 6 females of the two species.

| Replicate population | Selection Regime Te | Selection Regime Tu | Environment | Proportion of Te females (per replicate) |
|---|---|---|---|---|
| 1 | No Cadmium | No Cadmium | Cadmium | 0.9578 |
| 2 | No Cadmium | No Cadmium | Cadmium | 0.9992 |
| 3 | No Cadmium | No Cadmium | Cadmium | 0.9381 |
| 4 | No Cadmium | No Cadmium | Cadmium | 0.9622 |
| 5 | No Cadmium | No Cadmium | Cadmium | 0.9555 |
| 1 | No Cadmium | No Cadmium | No Cadmium | 0.6236 |
| 2 | No Cadmium | No Cadmium | No Cadmium | 0.6481 |
| 3 | No Cadmium | No Cadmium | No Cadmium | 0.7699 |
| 4 | No Cadmium | No Cadmium | No Cadmium | 0.7732 |
| 5 | No Cadmium | No Cadmium | No Cadmium | 0.7222 |
| 1 | Cadmium | No Cadmium | Cadmium | 0.9704 |
| 2 | Cadmium | No Cadmium | Cadmium | 0.9148 |
| 3 | Cadmium | No Cadmium | Cadmium | 0.9794 |

| | | | | |
|---|---|---|---|---|
| 4 | Cadmium | No Cadmium | Cadmium | 0.9567 |
| 5 | Cadmium | No Cadmium | Cadmium | 0.9571 |
| 1 | Cadmium | No Cadmium | No Cadmium | 0.6015 |
| 2 | Cadmium | No Cadmium | No Cadmium | 0.5636 |
| 3 | Cadmium | No Cadmium | No Cadmium | 0.9167 |
| 4 | Cadmium | No Cadmium | No Cadmium | 0.5065 |
| 5 | Cadmium | No Cadmium | No Cadmium | 0.7478 |
| 1 | No Cadmium | Cadmium | Cadmium | 0.9184 |
| 3 | No Cadmium | Cadmium | Cadmium | 0.8195 |
| 4 | No Cadmium | Cadmium | Cadmium | 0.9129 |
| 5 | No Cadmium | Cadmium | Cadmium | 0.9236 |
| 1 | No Cadmium | Cadmium | No Cadmium | 0.6484 |
| 3 | No Cadmium | Cadmium | No Cadmium | 0.6837 |
| 4 | No Cadmium | Cadmium | No Cadmium | 0.7885 |
| 5 | No Cadmium | Cadmium | No Cadmium | 0.8269 |
| 1 | Cadmium | Cadmium | Cadmium | 0.8336 |
| 3 | Cadmium | Cadmium | Cadmium | 0.9518 |
| 4 | Cadmium | Cadmium | Cadmium | 0.9070 |
| 5 | Cadmium | Cadmium | Cadmium | 0.9662 |
| 1 | Cadmium | Cadmium | No Cadmium | 0.7917 |
| 3 | Cadmium | Cadmium | No Cadmium | 0.7228 |
| 4 | Cadmium | Cadmium | No Cadmium | 0.8712 |
| 5 | Cadmium | Cadmium | No Cadmium | 0.8225 |
